# Unraveling genetic load dynamics during biological invasion: insights from two invasive insect species

**DOI:** 10.1101/2024.09.02.610743

**Authors:** Eric Lombaert, Aurélie Blin, Barbara Porro, Thomas Guillemaud, Julio S. Bernal, Gary Chang, Natalia Kirichenko, Thomas W. Sappington, Stefan Toepfer, Emeline Deleury

**Author notes:** **Corresponding author:** Eric Lombaert – Institut Sophia Agrobiotech – 400, route des chappes – BP 167 – 06903 Sophia-Antipolis Cedex – France.

## Abstract

Many invasive species undergo a significant reduction in genetic diversity, i.e. a genetic bottleneck, in the early stages of invasion. However, this reduction does not necessarily prevent them from achieving considerable ecological success and becoming highly efficient colonizers. Here we investigated the purge hypothesis, which suggests that demographic bottlenecks may facilitate conditions (e.g., increased homozygosity and inbreeding) under which natural selection can purge deleterious mutations, thereby reducing genetic load. We used a transcriptome-based exome capture protocol to identify thousands of SNPs in coding regions of native and invasive populations of two highly successful invasive insect species, the western corn rootworm (Chrysomelidae: *Diabrotica virgifera virgifera*) and the harlequin ladybird (Coccinelidae: *Harmonia axyridis*). We categorized and polarized SNPs to investigate changes in genetic load between invasive populations and their sources. Our results differed between species. In *D. virgifera virgifera*, although there was a general reduction in genetic diversity in invasive populations, including that associated with genetic load, we found no clear evidence for purging of genetic load, except marginally for highly deleterious mutations in one European population. Conversely, in *H. axyridis*, the reduction in genetic diversity was minimal, and we detected signs of genetic load fixation in invasive populations. These findings provide new insights into the evolution of genetic load during invasions, but do not offer a definitive answer to the purge hypothesis. Future research should include larger genomic datasets and a broader range of invasive species to further elucidate these dynamics.

## Introduction

Biological invasions represent a significant aspect of global change, profoundly impacting biodiversity through the alteration of species distributions worldwide, particularly in recent times due to the significant increase in human-assisted dispersal (Seebens et al. 2017). The key factors determining the success of invasive species are not yet fully understood, although numerous hypotheses have been proposed (Enders et al. 2018; Sherpa and Després 2021; Daly et al. 2023). Most proposals fail to account for one or more consistent characteristics of successful invasions: (i) the rarity of successful invasions resulting from introductions (Williamson and Fitter 1996), (ii) the lag time between initial introduction and invasion (Sakai et al. 2001), (iii) the frequent reduction in genetic diversity due to demographic bottlenecks (Nei et al. 1975), and (iv) the prevalence of multiple invasions originating from an initial invasive population (i.e. the bridgehead effect; Lombaert et al. 2010). These characteristics suggest that the success of invasions may stem from partially stochastic biological processes spanning multiple generations and combining both demographic and genetic mechanisms.

A hypothesis that aligns with these characteristics is that genetic load is purged in the initial stages of invasions, i.e. deleterious alleles that cause inbreeding depression are eliminated from the introduced population (Estoup et al. 2016). During the demographic bottleneck following introduction of a few individuals into a new environment, genetic drift intensifies and homozygosity increases. While genetic drift may randomly eliminate some deleterious alleles, thereby reducing part of the genetic load, it also contributes to the transformation of the masked load (i.e., the load which may become express in future generations; Bertorelle et al. 2022) into a realized load (i.e., the load which reduces fitness in the current generation; Bertorelle et al. 2022). This shift may result in a “mutational meltdown” (Lynch et al. 1995; Simberloff 2009), where expression of deleterious mutations increases risk of extinction, ultimately leading to failure of the introduced population to establish. Conversely, the increased homozygosity resulting from bottlenecks exposes recessive deleterious alleles to selection. In the short term, this exposure can reduce mean fitness, but if the population persists, purifying selection may progressively remove some highly deleterious alleles over multiple generations, potentially reducing genetic load (Dussex et al. 2023). Theoretically, such purges can occur under specific demographic (i.e., moderate reduction of population size) and genetic (i.e., highly deleterious, recessive alleles) conditions, optimizing exposure to natural selection (Crow 1970; Charlesworth et al. 1990; Glémin 2003; Robinson et al. 2023). The potential for such purging in introduced populations is particularly interesting for its role in facilitating successful invasions by reducing inbreeding depression and enabling inbred individuals to maintain high fitness levels.

Purging has been demonstrated empirically by measuring the evolution of various traits in artificially bottlenecked populations (Crnokrak and Barrett 2002; Avila et al. 2010). However, measuring life history traits can be challenging in the context of naturally occurring bottlenecks during biological invasions, and evidence of purging has been documented in only a few invasive species (Parisod et al. 2005; Mullarkey et al. 2013; Fountain et al. 2014; Marchini et al. 2016). One notable formal test of this hypothesis was conducted on the invasive Asian ladybird *Harmonia axyridis*, where measurement of life history traits revealed that invasive populations showed no evidence of the inbreeding depression observed in native ones, suggesting that deleterious alleles were purged during the invasion process (Facon et al. 2011). Overall, case studies focusing on the dynamics of genetic load during biological invasions have either examined a few life history traits (e.g. Facon et al. 2011) or used a single locus approach (e.g. Zayed et al. 2007), making it difficult to generalize the results.

Advances in population genomics over the past decade have provided promising avenues for investigating genetic load on large scales. Initially explored in humans (Lohmueller et al. 2008; Henn et al. 2015), these approaches have been applied in the fields of domestication (e.g. Schubert et al. 2014; Marsden et al. 2016; Makino et al. 2018; Wang et al. 2021) and conservation (Xue et al. 2015; Robinson et al. 2016; Grossen et al. 2020; Dussex et al. 2021; Ochoa and Gibbs 2021). These studies typically involve SNP-calling within coding regions, determination of ancestral SNP states, categorization of the severity of fitness reduction caused by derived alleles, and population comparisons while accounting for genetic drift using synonymous and/or intergenic polymorphisms. High quality genomic resources are essential for such studies, which may explain the limited application of these methods to invasive species, which are mostly non-model organisms. However, advances in genome sequencing technologies and bioinformatics have significantly reduced costs and improved accessibility, making these methods increasingly routine and affordable (Bertorelle et al. 2022).

In this study, we examined the dynamics of genetic load during the invasion of two successful insect species, namely the western corn rootworm, *Diabrotica virgifera virgifera*, and the harlequin ladybird, *Harmonia axyridis*, by directly measuring/estimating genetic load using genomic data from feral native and invasive populations. Importantly, our study was not designed to assess the instrumental role of purging in invasion success, but rather to investigate its occurrence during these invasions. A broader study of both successful and failed invasions across many species would be required to draw conclusions on purging’s role in invasion success. We used a pool-seq transcriptome-based exome capture protocol previously developed for non-model species (Deleury et al. 2020) to identify SNPs within coding sequences and categorize them as synonymous, moderately deleterious, or highly deleterious. Our results offer insights into the fate of the genetic load and provide valuable perspectives on the purge hypothesis in the context of biological invasions.

## Methods

### Species choice

The two species, *Diabrotica virgifera virgifera* (hereafter DVV) and *Harmonia axyridis* (HA), are good candidates for testing the purging of genetic load hypothesis. Both species are highly successful invaders with extensive invasive ranges (Gray et al. 2009; Roy et al. 2016). Additionally, their invasion routes are well-documented and supported by robust analyses using diverse datasets and methodologies (Miller et al. 2005; Ciosi et al. 2008; Lombaert et al. 2010, 2011, 2014, 2018). Both species exhibit bridgehead effects, meaning that certain invasive populations have played a crucial role in global dissemination (Guillemaud et al. 2011). Additionally, purging of genetic load was detected in HA in a laboratory study focusing on life-history traits (Facon et al. 2011).

### Design of exome capture probes for target enrichment

Design of the exome capture probes was performed as described in Deleury et al. (2020). We searched for peptide-coding sequences using FRAMEDP (v1.2.2; Gouzy et al. 2009) on the *de novo* transcriptomes described in Coates et al. (2021) and Vogel et al. (2017) for DVV and HA, respectively. BLASTX results (e-value ≤ 1e^-7^) of transcripts were used against the insect proteomes of *Tribolium castaneum, Anoplophora glabripennis, Dendroctonus ponderosae, Drosophila melanogaster* and the SwissProt database (v2016-02-17) for training. From the obtained coding sequences (CDS), we eliminated (i) *Wolbachia* and other putative endosymbiont sequences, (ii) CDS with > 1% missing nucleotide bases (Ns) or with more than four consecutive Ns, (iii) CDS with a GC% below 25 or above 75 and (iv) CDS with lengths < 400 bp or > 3500 bp. From the remaining 7,132 and 12,739 CDS for DVV and HA respectively, we drew at random c.a. 5.5 Mb for each species.

Probes based on the selected CDS were designed and manufactured by NimbleGen. In the case of DVV, repetitiveness of the probes was assessed based on the highest 15-mer frequency among the genomes of *Tribolium castaneum* (GCA_000002335.3), *Dendroctonus ponderosae* (GCA_000355655.1 and GCA_000346045.2) and DVV (the one available at the time, GCA_003013835.2). Probes with more than five close matches in the DVV genome were discarded. Close matches were defined as no more than five single-base insertions, deletions or substitutions using the SSAHA algorithm (Ning et al. 2001). Probes that matched sequences in the mitochondrial genome were also discarded. Random nucleotides were used to replace residual Ns in target sequences. For HA, the method used to ensure probe uniqueness is described in Deleury *et al*. (2020).

Overall, the final probe sets corresponded to a total of 4,151 CDS (5,282,603 bases across 12,017 regions of overlapping probes) and 5,717 CDS (5,347,461 bases across 6,400 regions of overlapping probes; Deleury et al. 2020) for DVV and HA respectively. This final set of probes was manufactured in the form of biotinylated DNA oligomers. One capture reaction contained 2,100,000 overlapping probes of 50 to 99 bp in length (mean length of 73.86 ± 4.46 bp for DVV, and 74.71 ± 4.92 bp for HA).

### Sample collection

For both species, the choice of populations to be sampled was based on previously known invasion routes (Figure S1; Miller et al. 2005; Lombaert et al. 2010, 2018), so that each invasive population could be compared to its source. We sampled adult DVV at 6 sites: two in the native area (Mexico), two in North America (Colorado and Pennsylvania, respectively corresponding to the core and front of the first invasive population; Lombaert et al. 2018), and two in Europe (Hungary, referred to the Central-Southeastern European population, and north-western Italy, which are independently derived from the same source area of eastern North America; Miller et al. 2005). Adult HA were sampled at four sites: two in the native area (Russia [Siberia] and China), and two in North America (Pennsylvania and Washington, corresponding to two independent outbreaks from the native area; Lombaert et al. 2010, 2014). None of the selected invasive populations resulted from multiple introductions. The east North American population of HA was previously hypothesized to be an admixture between two populations (Lombaert et al. 2011), but our analysis using ABC (approximate Bayesian computation; Beaumont et al. 2002) with synonymous SNPs from the current dataset indicates that this is unlikely (see Appendix S1 for details).

To determine the ancestral and derived alleles for each SNP (i.e. to polarize the alleles), outgroup species were also sampled. For DVV, we selected the closely related species *Diabrotica adelpha*, as well as the more phylogenetically distant chrysomelid *Cerotoma trifurcata* (Eben and de los Monteros 2013). In the case of HA, from phylogenetically closest to farthest (Tomaszewska et al. 2021), the sampled outgroup species were *Harmonia yedoensis, Harmonia conformis* and *Harmonia quadripunctata*. Complete information about samples is provided in Table 1 and Table S1.

**Table 1:** Description of *Diabrotica virgifera virgifera* (DVV) and *Harmonia axyridis* (HA) samples, including outgroup species (see Table S1 for BioSample accessions). Sources of invasive populations were determined based on known invasion routes (Figure S1; Miller et al. 2005; Lombaert et al. 2010, 2018), except for H-I-PEN for which we retraced invasion routes in this study (see Appendix S1 for details). Because the two native DVV populations are genetically almost indistinguishable, we selected D-N-MX1 as the source of D-I-COL based on the smallest pairwise *F*ST value between native and invasive populations (see Results section).

| Species | Population code name | Status (source) | Sampling site (Lat.; long.) | Sampling date | Haploid sample size |
| --- | --- | --- | --- | --- | --- |
| *Diabrotica virgifera virgifera* |  |  |  |  |  |
|  | D-N-MX1 | Native | Canatlán, Mexico (24.554; -104.741) | Oct. 2015 | 80 |
|  | D-N-MX2 | Native | Durango, Mexico (23.999; -104.614) | Oct. 2015 | 80 |
|  | D-I-COL | Invasive (D-N-MX1) | Fort Morgan, CO, USA (40.218; -103.869) | Aug. 2015 | 80 |
|  | D-I-PEN | Invasive (D-I-COL) | Landisville, PA, USA (40.119; -76.432) | Aug. 2015 | 80 |
|  | D-I-HUN | Invasive (D-I-PEN) | Kondoros, Hungary (46.736; 20.816) | July 2015 | 80 |
|  | D-I-ITA | Invasive (D-I-PEN) | Cuneo, Italy (44.463; 7.571) | July 2015 | 80 |
| *Harmonia axyridis* |  |  |  |  |  |
|  | H-N-CHI | Native | Beijing, China (40.057; 116.540) | Oct. 2015 | 72 |
|  | H-N-RUS | Native | Krasnoyarsk, Russia (55.992; 92.757) | Sept. 2015 | 72 |
|  | H-I-PEN | Invasive (H-N-CHI) | Landisville, PA, USA (40.119; -76.432) | Aug. 2015 | 72 |
|  | H-I-WAS | Invasive (H-N-CHI) | Spokane, WA, USA (47.665; -117.403) | Nov. 2015 | 72 |
| *Diabrotica adelpha* |  |  |  |  |  |
|  | D-OG-ADE | Outgroup DVV | Oaxaca State, Mexico (15.926; 97.151) | Jan. 2017 | 4 |
| *Cerotoma trifurcata* |  |  |  |  |  |
|  | D-OG-TRI | Outgroup DVV | Weldon, CA, USA (35.660; -118.330) | Oct. 2016 | 4 |
| *Harmonia yedoensis* |  |  |  |  |  |
|  | H-OG-YED | Outgroup HA | Beijing, China (40.031; 116.265) | Feb. 2015 | 2 |
| *Harmonia conformis* |  |  |  |  |  |
|  | H-OG-CON | Outgroup HA | Mouans-Sartoux, France (43.620; 6.934) | Oct. 2009 | 2 |
| *Harmonia quadripunctata* |  |  |  |  |  |
|  | H-OG-QUA | Outgroup HA | Rearing, Valbonne, France (NA) | July 2007 | 2 |

### DNA extraction, exome capture and pool sequencing

For each population, 4 legs from each of 40 individuals (DVV) or 2 legs from each of 36 individuals (HA) were pooled for DNA extraction with the Qiagen DNeasy Blood & Tissue kit, in accordance with the manufacturer’s recommendations. For the outgroup species, the same kit was used to extract DNA from a single individual each for the three *Harmonia* species, and from a pool of two individuals each for the *Diabrotica* and *Cerotoma* species.

Genomic libraries were prepared using NimbleGen SeqCap EZ HyperCap Library v2.0 and NimbleGen SeqCap EZ Library v5.0 for DVV and HA, respectively. In brief, for each of the population and outgroup samples, DNA (2 µg in 100 µl) was mechanically sheared to an average size of 200 bp using a Covaris S2 E210 device (6 cycles of 30 seconds each). In the case of DVV, the fragmented DNA was then divided into three technical replicates for each population. Subsequently, the fragments were subjected to end-repair, A-tailing and indexing (with one unique index per sample) using the KAPA Library Preparation kit designed for Illumina platforms. Following the ligation of Illumina adapters and indexes, only fragments falling within the size range of 250 to 450 bp were retained. A PCR amplification step consisting of 7 cycles was performed using standard Illumina paired-end primers, and the resulting amplicons were purified using AMPure XP beads (Beckman). The length, quality, and concentration of the prepared DNA fragments were assessed using a BioAnalyzer with Agilent High Sensitivity DNA Assay, along with a Qubit.

For each capture (five for DVV and two for HA), we used a total of approximately 1 µg of amplified DNA for exome enrichment, combining multiple samples in proportions that allowed for equimolarity between population samples (see Table S2 and Table S3). This enrichment was performed using the capture probes described above, following the guidelines of either the NimbleGen SeqCap EZ HyperCap Library Protocol v2.0 or the SeqCap EZ Library Protocol v5.0. Following each capture, we conducted two parallel PCRs, each comprising 14 cycles, on the elution solution. In the case of HA, the resulting PCR products were combined. Subsequently, all PCR products underwent purification using AMPure XP beads. The length, quality, and concentration of the final DNA fragments were assessed using a BioAnalyzer equipped with the Agilent High Sensitivity DNA Assay and a Qubit fluorometer. For sequencing, we used one lane of an Illumina HiSeq3000 sequencer per species, following the manufacturer’s instructions, in paired-end mode for 150 cycles. After sequencing, the data were demultiplexed and exported as FastQ files, and the libraries were processed independently. For DVV, the FastQ files of technical replicates were merged prior to subsequent analysis.

### Mapping, SNP calling and annotation

Sequence quality assessment was conducted using FastQC v0.11.5 (Andrews 2010). Subsequently, adapter sequences were removed, and low-quality base pairs were eliminated using *Trimmomatic* v0.35 (Bolger et al. 2014), with the following parameter settings: ILLUMINACLIP:TruSeq-file.fa:2:30:10 LEADING:25 TRAILING:25 SLIDINGWINDOW:5:30 MINLEN:75. We used almost the same parameters for the outgroup species sequences, but with less stringency on quality of reads (SLIDINGWINDOW:5:20).

Filtered reads were mapped onto the genome assemblies PGI_DIABVI_V3a (GenBank identifier: GCA_917563875.2) and icHarAxyr1.1 (GenBank identifier: GCA_914767665.1), for DVV and HA respectively, using default options of the *bwa-mem* aligner v0.7.15 (Li 2013). We used the *SAMtools* v1.15.1 software package (Li et al. 2009) and its *fixmate, view*, and *markdup* tools to perform the following operations sequentially: (1) removal of unmapped reads and secondary alignments, (2) filtering out read alignments with a mapping quality Phred-score <20 and improperly paired reads, and (3) identification and removal of PCR duplicates. Processing filtered reads for the outgroup species followed the same steps, including mapping to the genome of the corresponding focal species, except that PCR duplicates were not removed.

For each species, we conducted variant calling on the resulting *bam* alignment files using Freebayes v1.3.6 software (Garrison and Marth 2012), with a focus on exonic regions (options *-t* exons.bed *--pooled-continuous --min-alternate-count* 1 *--min-alternate-fraction* 0.001 *--min-alternate-total* 1 *--min-coverage* 80 *--use-best-n-alleles* 3). The resulting *vcf* file was subsequently filtered using the *view* tool within *bcftools* v1.13 software (Danecek et al. 2021) and an in-house script to retain only true bi-allelic SNPs. Annotation was then performed with the *SnpEff* program v5.0 (Cingolani et al. 2012b).

### Allele polarization

SNP positions were extracted from both *vcf* files using the bcftools query tool. Subsequently, for each outgroup species, we identified the nucleotides (or absence of typing) at positions matching those in their respective focal species using the *samtools mpileup* tool on the previously generated *bam* alignment files.

To polarize the SNPs, we used *est-sfs* software v2.03 (Keightley and Jackson 2018) with the Kimura 2-parameter model. The input files contained data from one native population of the focal species, together with the corresponding outgroup species, ensuring a consistent phylogenetic topology for the software. Given that *est-sfs* computes the probability of the most frequent allele being ancestral solely for polymorphic SNPs within the native population, we used probabilities from a parallel analysis for monomorphic loci, incorporating an extra haplotype for each allele. The complete process was repeated twice for each focal species, considering the availability of two native populations for each. This resulted in two *est-sfs* probabilities per focal species, both reflecting the likelihood of the most frequent allele being the ancestral one.

To consider a SNP as polarized, we applied the following rules. If both probabilities were above 0.5 for the same allele, and at least one of the probabilities exceeded 0.75, we considered the most frequent allele as the ancestral one. If both probabilities were below 0.5 for the same allele, with at least one below 0.25, and if there were no more than two distinct nucleotides present in total across the focal and outgroup species combined, we considered the less frequent allele to be the ancestral one. SNPs that did not meet these criteria were considered as not polarized but were retained for computations that did not require polarization. Note that, because the est-sfs probabilities were predominantly close to 1 (Figure S2), the threshold value had minimal impact on the number of polarized SNPs (Figure S3).

### Additional filters and categorization of deleterious mutations

Key information was extracted from the *vcf* file using the *SnpSift* program v5.0 (Cingolani et al. 2012a), including SNP coordinates, reference and alternative alleles, allele depths, predicted effects, impact annotations, and protein-level changes. Additional filtering was performed with an in-house R script (R Core Team 2021). First, we retained only biallelic SNPs with coverage greater than 50 reads, falling below the 95th percentile of overall coverage in each pool, and exhibiting a minor allele frequency exceeding 0.01 in at least one population. Second, we retained SNPs located in coding regions with unambiguous annotations. Finally, we excluded SNPs located on the X-chromosome in the specific case of HA, for which this information was available.

*SnpEff* annotations were used to categorize the severity of fitness loss due to mutations. Synonymous variants were used as proxies for neutral polymorphism. Non-synonymous mutations were categorized as either “missense” (considered as potentially moderately deleterious, involving amino acid changes) or “LoF” (loss-of-function, considered as potentially highly deleterious, involving gain or loss of stop codons).

### Genetic diversity and genetic load analyses

We computed several descriptive statistics on the whole set of SNPs. At the inter-population level, we assessed genetic differentiation by calculating pairwise *F*_ST_ values using the R package *poolfstat* v2.1.1 (Gautier et al. 2022). At the intra-population level, we used in-house R scripts to compute the synonymous expected heterozygosity *H*_*eS*_, as a measure of diversity, and the ratio of non-synonymous to synonymous expected heterozygosity *H*_*eN*_/*H*_*eS*_, which provides an indirect measure of the efficacy of selection. We used a 100-block jackknife resampling approach to estimate means and standard errors. Additionally, we estimated bottleneck intensities for all invasive populations of both species using ABC analyses (See Appendix S1 for details).

Using polarized SNPs, we reported for each population and each severity category the proportion of positions for which the derived allele is absent (frequency *f* = 0), rare (*f* > 0 and *f* ≤ 0.1), common (*f* > 0.1 and *f* < 1), or fixed (*f* = 1). The proportion of rare derived alleles serves as a proxy for the masked load, whereas common and fixed alleles serve as a proxy for the realized load. Additionally, within each severity category and population, we calculated the mean derived allele frequency and estimated standard errors via a 100-block jackknife resampling approach. Within each species and severity category, mean derived allele frequencies were compared among populations using z-tests with false discovery rate (FDR; Benjamini and Hochberg 1995) correction for multiple comparisons.

Finally, polarized SNPs were used to compute the *R*_*XY*_ statistic, which allows assessment of relative excess or deficit of derived alleles within specific categories of deleterious mutations in one population compared to another (Appendix S2; Do et al. 2015; Xue et al. 2015). This statistic effectively corrects for variation due to demography using neutral mutations (synonymous SNPs in our case) as a reference point. In our analyses, population *X* is an invasive population, and population *Y* is its source population (provided in Table 1). An *R*_*XY*_ value of 1 indicates an equivalent relative genetic load in both populations, whereas values greater or less than 1 suggest an excess (including fixation) or a deficit (including purge) of the genetic load in the invasive population *X*, respectively. We assessed the significance of any deviation from 1 using a z-score two-tailed test based on the 100-block jackknife resampling method.

## Results

### SNP-calling and genomic variation

In the target enrichment experiments, we obtained mean numbers of raw read sequences of 50,196,834 and 63,360,676 for DVV and HA, respectively (BioProject PRJNA1079689; Table S1). After trimming, 85.0% and 93.8% of the sequences were retained for DVV and HA respectively. For outgroup species, the total number of raw reads was highly variable, with a mean of 2,631,775 of which 97.8% were conserved after trimming (see detailed information for all libraries, including outgroup species, in Table S4). Mapping, calling and filtering identified 66,274 SNPs (of which 62,034 could be polarized) and 169,755 SNPs (of which 169,102 could be polarized) within coding sequences for DVV and HA respectively (see Table S5 for details).

In the case of DVV, pairwise *F*_ST_ computed on the full set of SNPs showed patterns very close to those expected from results of previous analyses of microsatellite datasets (Figure S4). The differentiation between the two native Mexican populations was almost negligible with a mean *F*_ST_ estimate below 0.001. Given that the *F*_ST_ estimates between the invasive populations and the native sample D-N-MX1 (mean *F*_ST_ of 0.121) were consistently lower than those between the invasive populations and D-N-MX2 sample (mean *F*_ST_ of 0.128), D-N-MX1 was therefore considered the source of the D-I-COL sample (see Table 1). The Pennsylvania and Colorado populations had a low pairwise *F*_ST_ of 0.012, consistent with an Eastern expansion from Colorado (Figure S1), with plausible continuous gene flow. Conversely, all other *F*_ST_ estimates were relatively high, particularly between the native and European populations (mean *F*_ST_ value = 0.152). The highest value was observed between the two European populations at 0.169, confirming previous results indicating these populations originated from two independent introductions (Miller et al. 2005).

In the case of HA, *F*_ST_ estimates were moderate and fairly homogeneous, ranging from 0.020 to 0.050 (Figure S4). The smallest values were found between the eastern native population H-N-CHI and both invasive populations. The *F*_ST_ between invasive populations was slightly larger, consistent with previous results indicating independent origins for the two North American outbreaks (Lombaert et al. 2010). The highest values were observed between the western native population H-N-RUS and all other populations.

Within each species, invasive populations were characterized by reduced synonymous heterozygosity compared to their native counterparts, reflecting demographic bottlenecks encountered during the invasion process (Figure 1, x axis). One exception was the western native population of HA (H-N-RUS), which exhibited the lowest diversity within this species. Overall, the loss of diversity was more pronounced in DVV invasive populations, consistent with our ABC results, which indicate that bottlenecks were generally more intense in DVV than in HA (see Appendix S1). The ratio of non-synonymous to synonymous expected heterozygosity *H*_*eN*_/*H*_*eS*_ is typically expected to increase in populations experiencing pronounced drift, as the reduced efficacy of selection allows more non-synonymous mutations, including potentially deleterious ones, to persist. In line with this expectation, we observed a higher ratio in invasive populations (Figure 1, y axis), and a negative correlation between this ratio and synonymous expected heterozygosity within each species (Pearson’s *r* = -0.95, *P* < 10^-2^ for DVV; Pearson’s *r* = -0.90, *P* < 10^-1^ for HA; Figure 1). Overall, the differences between native and invasive populations were more pronounced in DVV, with invasive ranges showing lower diversities and higher *H*_*eN*_/*H*_*eS*_ ratios. This could suggest a reduced efficacy of selection, although other factors may also contribute to the observed patterns.

**Figure 1:**
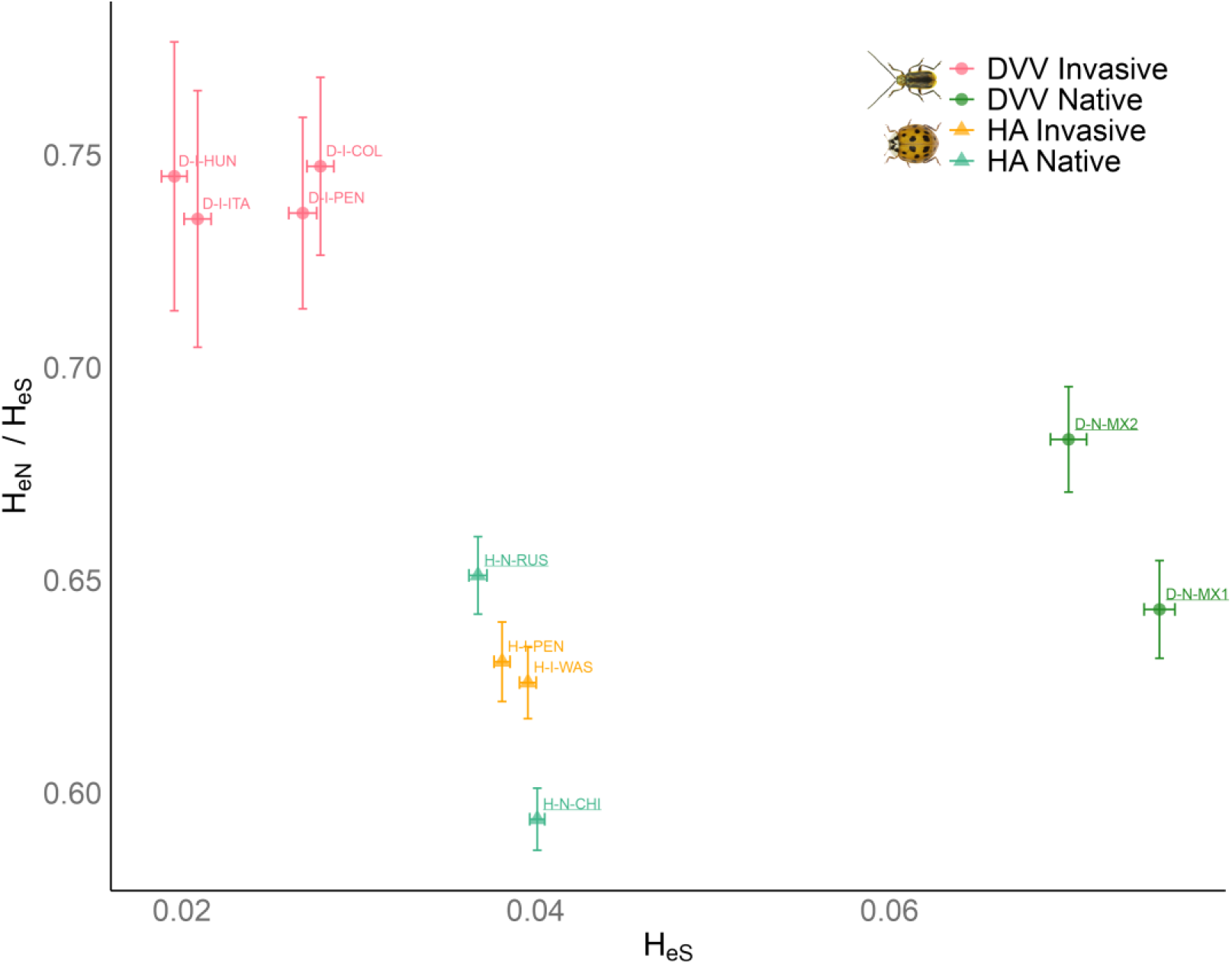
Ratio of non-synonymous to synonymous expected heterozygosity *HeN*/*HeS* vs. the synonymous expected heterozygosity *HeS* of each *Diabrotica virgifera virgifera* (DVV) and *Harmonia axyridis* (HA) population. Means and standard errors (error bars) were determined from 100-block jackknife resampling. See Table 1 for population codes (underlined population codes correspond to the native populations).

### Genetic load on polarized SNPs

In all populations studied and for each species, derived alleles were mostly rare (with frequencies below 0.1) or absent (Figure 2). The highest derived allele frequencies consistently showed a higher prevalence in synonymous positions compared to non-synonymous positions. In DVV, we observed a substantial loss of non-synonymous derived alleles in invasive populations compared to native ones. Although this was characterized by a reduction in both masked load (proportion of non-synonymous SNPs with rare derived alleles ranging from 0.496 to 0.523 in native populations and from 0.145 to 0.188 in invasive populations) and realized load (proportion of non-synonymous SNPs with frequent or fixed derived alleles ranging from 0.075 to 0.077 in native populations and from 0.046 to 0.051 in invasive populations), it also came with slightly higher proportions of fixed derived alleles. This trend was less pronounced for mutations inferred to be highly deleterious (LoF) than for those inferred to be moderately deleterious (missense). For HA, the disparities between native and invasive populations were considerably less pronounced (Figure 2). Compared to their native Chinese source, HA invasive populations displayed a slight reduction in the proxy for masked load (proportion of non-synonymous SNPs with rare derived alleles: 0.455 in the Chinese population, and 0.347-0.381 in invasive populations) and a slight increase in the proxy for realized load (proportion of non-synonymous SNPs with frequent or fixed derived alleles: 0.019 in the Chinese population, and 0.024-0.027 in invasive populations). Again, this trend was less pronounced for highly deleterious mutations (Figure 2).

**Figure 2:**
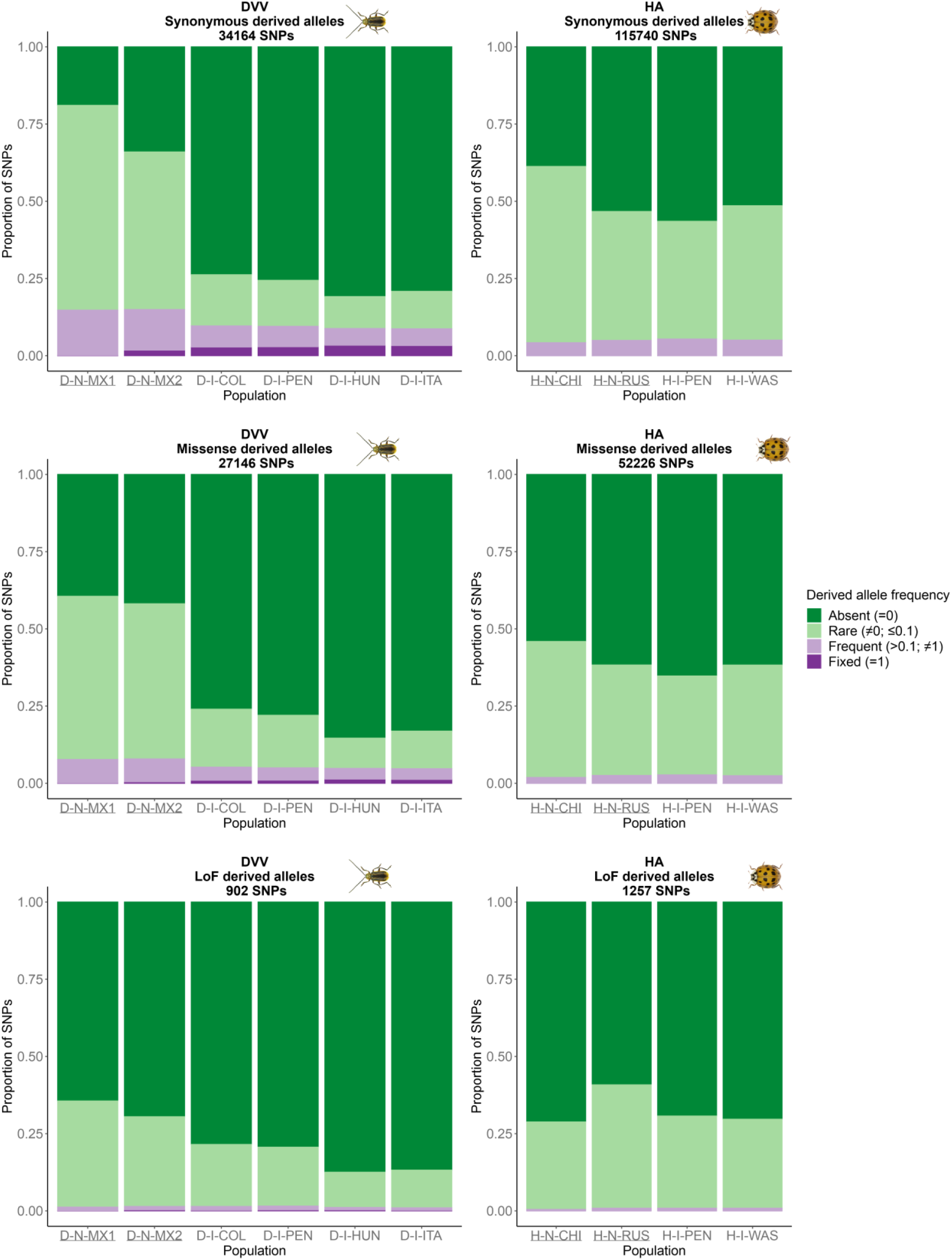
Proportion of derived alleles found in each population of *Diabrotica virgifera virgifera* (DVV) and *Harmonia axyridis* (HA) within categories of putative severity, classified by their frequency. See Table 1 for population codes (underlined population codes correspond to the native populations).

As expected, the mean derived allele frequencies within populations decline with increasing putative severity in both species (Figure 3). For DVV, synonymous and missense mean derived allele frequencies were consistently significantly lower in the invasive populations than in the native populations (*P* < 10^-4^ for all comparisons after FDR correction), whereas no significant differences were observed between native populations or between invasive populations (*P* > 0.3 for all comparisons after FDR correction). Within each of the three severity categories, mean derived allele frequencies were virtually identical in the four HA populations (P > 0.02 for all comparisons after FDR correction). No differences in LoF derived allele frequencies were significant (*P* > 0.2 for all comparisons after FDR correction), for either species. Overall, derived allele frequencies were lower in HA than in DVV.

**Figure 3:**
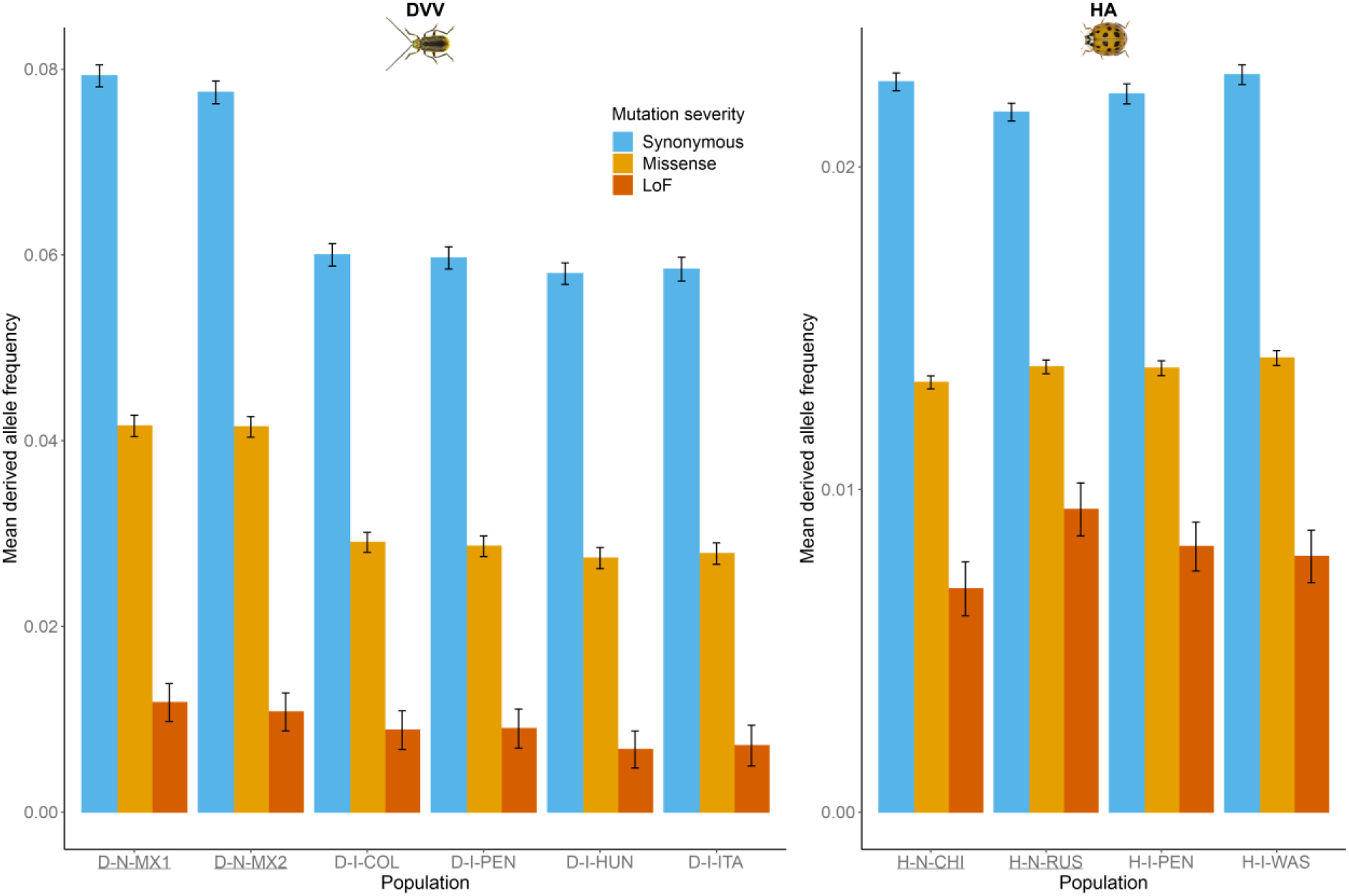
Mean derived allele frequencies in all *Diabrotica virgifera virgifera* (DVV) and *Harmonia axyridis* (HA) populations in all three severity categories. Means and standard errors (error bars) were determined from 100-block jackknife resampling. See Table 1 for population codes (underlined population codes correspond to the native populations).

Finally, the *R*_*XY*_ ratio revealed no significant differences in relative frequencies of missense derived alleles between invasive populations and their respective sources in DVV (Figure 4). For putatively highly deleterious loss-of-function alleles, *R*_*XY*_ values sometimes deviate sharply from 1, but the number of SNPs is small (Table S5), and most differences are not statistically significant. The only exception is found in the population from Hungary (D-I-HUN), which shows reduced relative frequencies within the putatively highly deleterious category (LoF) of alleles (Figure 4). Regarding HA, slightly higher loads were significant in three of four comparisons across severity categories between the two invasive populations (H-I-PEN and H-I-WAS) and the native Chinese population (Figure 4).

**Figure 4:**
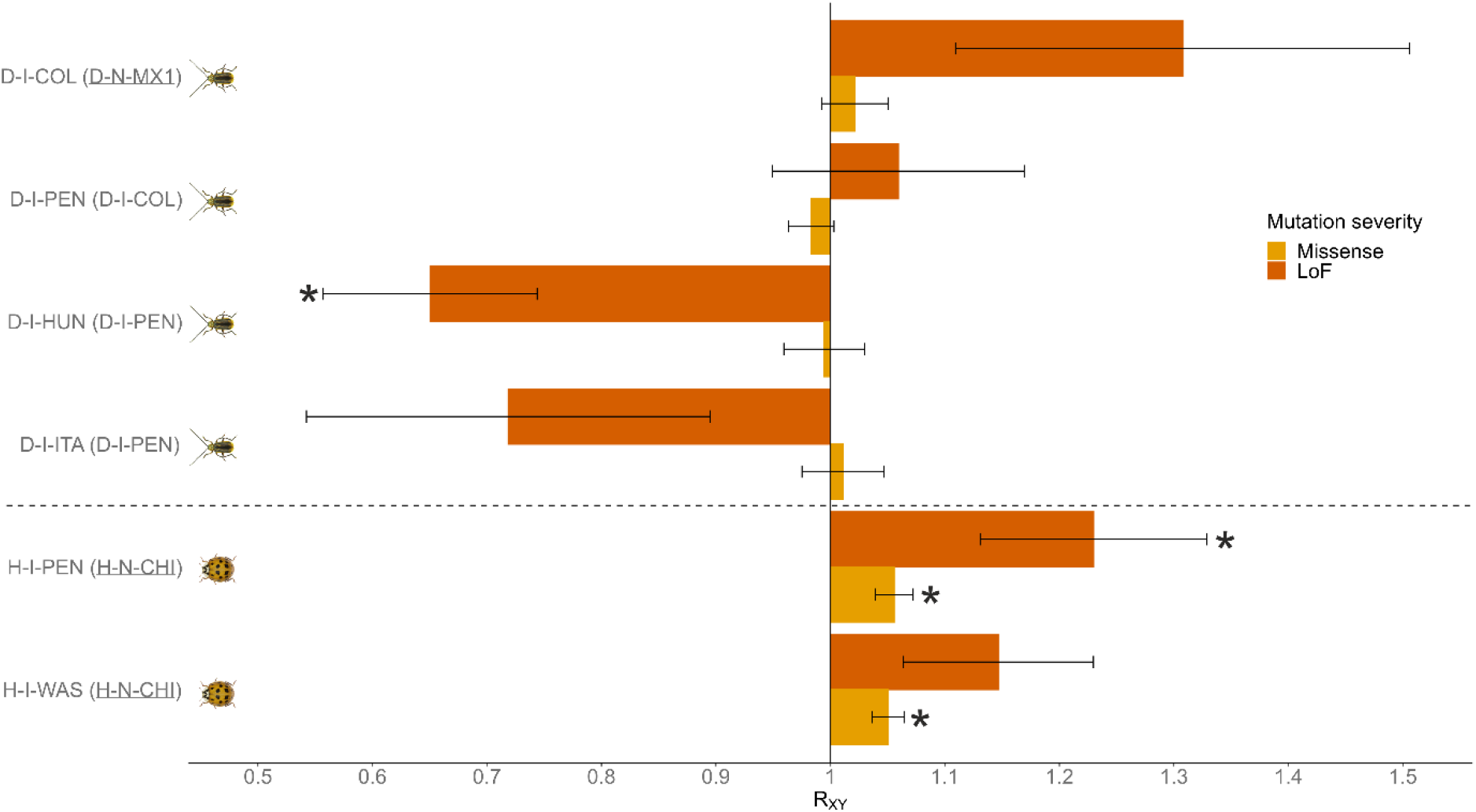
Relative mutation load estimated as the *RXY* ratio of derived alleles for moderate (missense) and high (LoF) severity categories in populations of *Diabrotica virgifera virgifera* (D) and *Harmonia axyridis* (H). A ratio below or above 1 indicates, respectively, a relative frequency deficit or excess of derived alleles of indicated severity category in the invasive population (listed first) compared to its population source (in parentheses). Means and standard errors (error bars) were determined from 100-block jackknife resampling. A star indicates that the corresponding *RXY* value is significantly different from 1 (z-score two-tailed test). For population codes, refer to Table 1 (underlined population codes correspond to native populations).

## Discussion

Exploring the evolution of genetic load using genomic data has long been restricted to model species. Here, we successfully identified putatively deleterious derived alleles within coding regions in two non-model invasive insect species using a pool-seq transcriptome-based exome capture protocol (Deleury et al. 2020). By comparing native and invasive populations, we tested the hypothesis that genetic load was purged during the invasion process, which may enhance invasion success of certain populations (Sherpa and Després 2021; Daly et al. 2023). Given that bottlenecks amplify genetic drift and weaken selection effectiveness (Crow 1970; Glémin et al. 2003), the likelihood of purging is expected to increase for recessive, highly deleterious mutations, while moderately deleterious mutations may still experience some degree of drift, leading to higher frequencies (Agrawal and Whitlock 2011). Our study exploring the dynamics of the genetic load in DVV and HA yielded mixed results, underscoring the nuanced interplay of evolutionary forces in invasive species.

### *Evolution of the genetic load in* Diabrotica virgifera virgifera

DVV exhibits marked genetic differences between native and invasive populations. Consistent with previous studies (Ciosi et al. 2008; Lombaert et al. 2018), invasive populations, including the long-established population in Colorado, display sharp declines in genetic diversity, consistent with the strong bottlenecks inferred from ABC analyses. Compared to their native counterparts, all invasive populations generally exhibit lower frequencies of derived alleles, regardless of their potential fitness impacts, thus confirming the substantial role of genetic drift in the invasion process of the species. While this overall reduction in genetic load within invasive populations is apparent, it is worth emphasizing the presence of a slightly larger fixed load.

In North America, despite a strong initial loss of genetic diversity, invasive populations showed no significant relative deficit or excess of deleterious mutations compared to their source, although a trend toward an excess of loss-of-function variants was observed.

Interestingly, the oldest European outbreak in Central-Southeastern Europe (here a sample from Hungary) exhibited a significant deficit of “loss-of-function” deleterious mutations compared to its source population, while “missense” deleterious mutations did not differ significantly. This suggests that the demographic and selective constraints in this population were effective at reducing highly deleterious mutations through a combination of genetic drift and purifying selection, while moderately deleterious mutations have been less affected. Such differences in evolutionary trajectories according to mutation severity are theoretically expected (Whitlock 2002; Caballero et al. 2017; Dussex et al. 2023) and have been observed in other species experiencing severe bottlenecks (Xue et al. 2015; Khan et al. 2021; Ochoa and Gibbs 2021; Wang et al. 2023).

Finally, the invasive northwestern Italian population, despite originating from an independent introduction (Miller et al. 2005), showed patterns similar to that of the Central-Southeastern European population. However, it did not exhibit significant signals of deficit or excess of deleterious alleles, regardless of the mutation severity considered.

### *Evolution of the genetic load in* Harmonia axyridis

In HA, the overall loss of genetic diversity in both invasive populations was relatively low, in agreement with previous findings (Lombaert et al. 2010, 2011) and consistent with our ABC results suggesting that bottlenecks were relatively mild in HA invasive populations. While the putative masked genetic load decreased slightly, the putative realized load, although consistently low in all populations, showed no sign of decreasing and even exhibited a slight increase. This observation aligns with a significant trend, although subtle, towards a relative excess of deleterious mutations in invasive populations compared to their native source, regardless of putative fitness impact. This tendency to fix alleles contributing to genetic load, likely driven by drift, is significant across all populations and severity categories, except for the highly deleterious mutations in western North America (sampled in Washington).

These findings diverge from a previous study based on life history traits, which demonstrated that invasive populations of HA experienced minimal inbreeding depression while maintaining fitness comparable to outbred native populations (Facon et al. 2011). That study has greatly contributed to the hypothesis of purging genetic load as a key mechanism driving successful biological invasions (Estoup et al. 2016; Sherpa and Després 2021; Daly et al. 2023). Given the markedly different methodologies used in the study by Facon et al. (2011) and our study, it is challenging to ascertain why our results do not align. It is conceivable that a few key genes could have disproportionate effects on invasion success, an aspect difficult to quantify using our approach, particularly in the context of the overall low genetic load previously described in this species (Tayeh et al. 2013). This issue is further complicated by the imperfect nature of our categorization, where missense mutations likely cover a broad spectrum of severity levels, potentially obscuring the influence of key genes. Moreover, very highly deleterious mutations, which are critical contributors to inbreeding depression, are expected to segregate at extremely low frequencies in large populations and may thus escape detection in our sampling due to their rarity. Finally, our exome capture protocol only targets a fraction of the exome, approximately one-third, and a broader approach may yield qualitatively different results.

### Conclusions, limits and perspectives

We tested the hypothesis that genetic load was purged during the invasion of two insect species, a crop pest (DVV) and a predator (HA). At first glance, our results offer a nuanced perspective. In the case of DVV, we were unable to detect any significant evolution of the genetic load, except for the purge of highly deleterious mutations in the invasive Central-Southeastern European population. For HA, we observed subtle evidence suggesting a tendency toward fixation, rather than toward purge, of the genetic load. Although these results are not entirely contradictory, they may stem from differences in initial genetic composition, varying demographic history, and divergent ecological niches. For instance, the two species experienced markedly different bottleneck intensities, yet our data do not allow us to establish a direct link between these differences and the observed dynamics of genetic load.

While population genomics approaches to assess the evolution of genetic load have become increasingly popular in conservation and domestication biology (Moyers et al. 2018; Bertorelle et al. 2022), our study is, to our knowledge, the first to use these methods to specifically investigate the evolution of genetic load in the context of biological invasions. However, it does not yet provide a comprehensive and definitive answer to the question addressed. One limitation lies in our reliance on allelic frequencies derived from pool-seq data, which precludes access to genotypic frequencies and therefore limits our ability to precisely estimate inbreeding depression and explore the non-linear effects of genetic load (Bataillon and Kirkpatrick 2000). While we used complementary statistics to mitigate this issue, future studies based on individual-level sequencing would allow for more accurate assessments of genetic variation and its relationship to genetic load. Additionally, to further broaden our findings and enhance our statistical power, particularly for highly deleterious mutations, it will be essential to consider the entire exome rather than only a fraction. Another promising avenue involves categorizing mutation severity by quantifying evolutionary constraints within a phylogenetic framework, which could enhance our understanding of genetic load evolution, including in non-coding regions (Davydov et al. 2010). Furthermore, extending this research to a wider range of species would enhance our ability to identify potential universal patterns in the evolution of genetic load during biological invasions. Finally, to test not only the occurrence of purging in invasions but also its role in the success of invasions, we must compare the load dynamics in introductions that fail with those that succeed – a comparison that is rarely feasible (Zenni and Nuñez 2013), except in controlled invasions such as in classical biological control (Fauvergue et al. 2012).

## Acknowledgments

We thank our colleagues Pamela Bruno, Charlyne Martignier-Jaccard, Pascal Maignet and Wang Su for *Diabrotica adelpha, Harmonia quadripunctata* and *Harmonia yedoensis* samples. We also thank Emmanuelle Murciano-Germain for administrative assistance. Sequencing was performed at the GENOTOUL GeT platform (https://get.genotoul.fr/en/).

## Data availability

Target capture pool sequencing files and all sequencing data have been deposited in Sequence Read Archive, National Center for Biotechnology Information, under project PRJNA1079689 (see also Table S1): https://www.ncbi.nlm.nih.gov/bioproject/PRJNA1079689. All scripts used in this study have been deposited at Data INRAE: https://doi.org/10.57745/ESQFDB

## Author contributions

EL and ED designed the study. JB, GC, NK, TS and ST managed the collection of samples. AB and ED prepared the samples and the libraries. ED and BP developed the bioinformatics pipelines. EL and ED analyzed the data. EL and TG wrote the paper. All authors have revised and approved the final manuscript.

