## Supplementary material for "Unraveling genetic load dynamics during biological invasion: insights from two invasive insect species"

| | |
|---|---|
| *Appendix S1* | *p. 2-9* |
| *Appendix S2* | *p. 10* |
| *Figure S1* | *p. 11* |
| *Figure S2* | *p. 12* |
| *Figure S3* | *p. 13* |
| *Figure S4* | *p. 14* |
| *Table S1* | *p. 15* |
| *Table S2* | *p. 16* |
| *Table S3* | *p. 16* |
| *Table S4* | *p. 17* |
| *Table S5* | *p.18* |
| *Supporting information references* | *p. 19-20* |

### Appendix S1
### Approximate Bayesian analyses
### (origin of the eastern North American population of *Harmonia axyridis*; estimation of bottleneck intensities)

#### 1. ABC analyses: general methodology

Approximate Bayesian computation analyses (ABC; Beaumont et al. 2002) were conducted to (i) infer the invasion history of the eastern North American population of *Harmonia axyridis* and (ii) estimate the bottleneck intensities of all invasive populations for both species, *Harmonia axyridis* (two populations) and *Diabrotica virgifera virgifera* (four populations). ABC is a model-based Bayesian method that computes posterior probabilities of historical scenarios and parameter estimation using massive simulations of genetic data. All simulations and ABC analyses were performed with DIYABC-RF software (Collin et al. 2021), which integrates diyabc v1.1.54 with abcranger v1.16.69. Gene flow was not modeled in these simulations, as it is not supported by DIYABC-RF, but the software remains well-suited for Pool-Seq data and large-scale invasions.

The pool-datasets we used all consisted of 10,000 randomly selected synonymous SNPs, each of which met a minimum read count of 4 across all samples. We simulated 10,000 datasets per scenario with diyabc v1.1.54.

In all analyses, simulated and observed datasets were summarized using the complete set of summary statistics proposed by DIYABC-RF for SNP markers:

- Proportion of monomorphic loci for each population, as well as for each pair, triplet and quadruplet of populations;
- Heterozygosity for each population and for each pair of populations (mean and variance over loci; Hivert et al. 2018);
- $F_{ST}$-related statistics for each population (i.e., population-specific FST; Weir and Goudet 2017), for each pair, triplet, quadruplet and overall populations when the dataset includes more than four populations (mean and variance over loci; Hivert et al. 2018);
- Allele shared Patterson's f-statistics for each triplet (f3-statistics) and quadruplet (f4-statistics) of populations (Patterson et al. 2012; mean and variance over loci; Leblois et al. 2018);
- Nei's (1972) distance for each pair of populations (mean and variance over loci);
- Maximum likelihood coefficient of admixture computed for each triplet of populations (mean and variance over loci; adapted from Choisy et al. 2004).

#### 2. Origin of the eastern North American population of *Harmonia axyridis*

##### *2.1 Background and motivation*

The first invasive population of *Harmonia axyridis* was observed in Louisiana state in 1988 (Chapin and Brou 1991). Since then, this population has expanded across much of the eastern USA and Canada (Lombaert et al. 2014). The origin of this population was investigated using microsatellite data (Lombaert et al. 2011), and the results suggested that it originated from an admixture between the two main genetic clusters found in the native area: the eastern cluster (China, Japan, eastern Siberia) and the western cluster (western Siberia, Kazakhstan). Recently, several unpublished pieces of evidence obtained from more numerous markers and more powerful analysis methods have suggested that this result might be incorrect, and that this invasive population might instead originate solely from the eastern native cluster. Therefore, we decided to use an ABC framework to infer the native origin of the eastern North American invasive population with the new dataset available in the present study.

##### *2.2 ABC analyses: posterior probabilities of historical scenarios*

In our competing scenarios, the eastern North American invasive population (represented by the H-I-PEN sample; see Table 1) originated from either (1) the east of the native area (represented by the H-N-CHI sample), (2) the west of the native area (represented by the H-N-RUS sample), or (3) an admixture of both.

The scenarios were coded as follows (see Figure A1 for graphical representations):

Scenario 1:
```
Ne Nw Nlou Ngh
0 sample 1
0 sample 2
0 sample 3
tlou-DBlou VarNe 3 NFlou
tlou VarNe 3 Ne
tge merge 1 3
tne merge 4 1
tnw merge 4 2
```

Scenario 2:
```
Ne Nw Nlou Ngh
0 sample 1
0 sample 2
0 sample 3
tlou-DBlou VarNe 3 NFlou
tlou VarNe 3 Nw
tgw merge 2 3
tne merge 4 1
tnw merge 4 2
```

Scenario 3:
```
Ne Nw Nlou Ngh Ne Nw
0 sample 1
0 sample 2
0 sample 3
tlou-DBlou VarNe 3 NFlou
tlou split 3 5 6 r
tge merge 1 5
tgw merge 2 6
tne merge 4 1
tnw merge 4 2
```

Population numbers are as follows: 1 for East Asia (effective population size $N_e$; sample H-N-CHI); 2 for West Asia (effective population size $N_w$; sample H-N-RUS); 3 for eastern North America (effective population size $N_{lou}$; sample H-I-PEN); 4 for an unsampled ancestral native population (effective population size $N_{gh}$). The bottleneck lasts $DB_{lou}$ generations, and the effective number of founders during this duration is $NF_{lou}$. The eastern North American population merges into an unsampled native population at $t_{lou}$. Eastern and western unsampled native populations merge into their corresponding native population at $t_{ge}$ and $t_{gw}$ respectively. Populations 1 and 2 merge into the ancestral native population at $t_{ne}$ and $t_{nw}$, respectively. Parameter $r$ is an admixture rate.

Historical and demographic parameter values for simulations were drawn from prior distributions defined as follows:
```
NFlou N LU[2,1000,0,0]
Ne N UN[100,100000,0,0]
Nw N UN[100,100000,0,0]
Nlou N UN[100,100000,0,0]
Ngh N LU[100,20000,0,0]
tlou T UN[68,73,0,0]
DBlou T UN[1,10,0,0]
tge T LU[68,3000,0,0]
tgw T LU[68,3000,0,0]
tne T LU[100,3000,0,0]
tnw T LU[100,3000,0,0]
r A UN[0.1,0.9,0,0]
```
With the following conditions:
```
tgw>=tlou
tge>=tlou
tnw>=tgw
tne>=tge
```

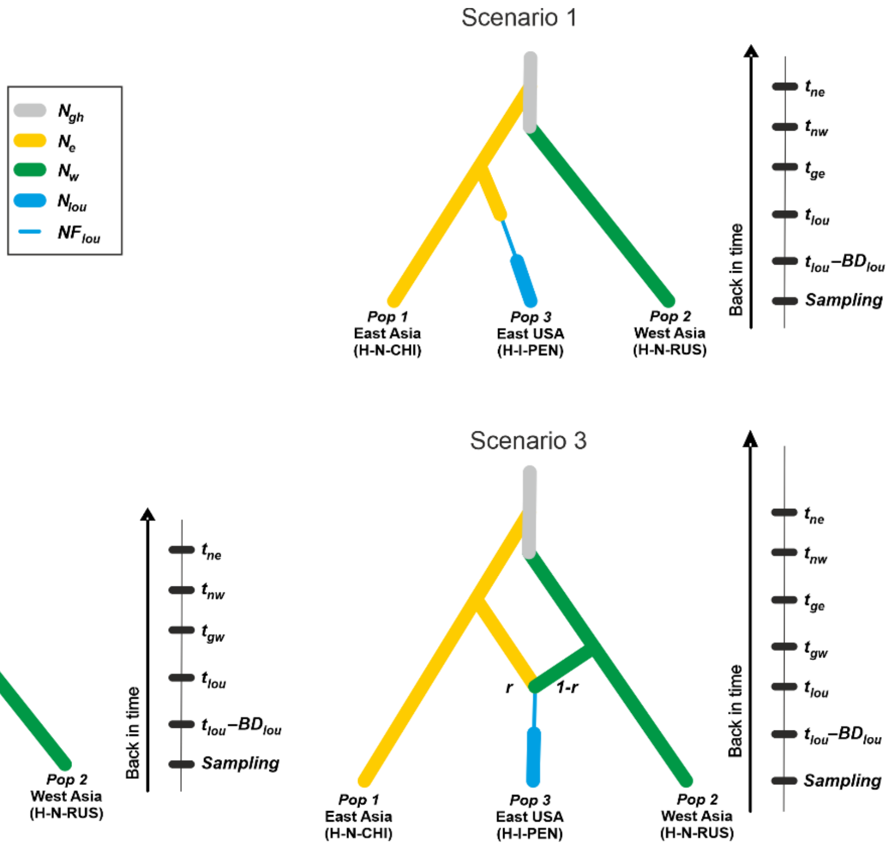


**Figure A1**: Schematic representation of each set of scenarios used in the ABC analyses to decipher the origin of the eastern North American population of *Harmonia axyridis*. Time is not to scale.

Since the dataset includes three populations, the total number of summary statistics was 50 (plus two LDA axes).

To compare the scenarios, we used a random forest procedure (RF; Breiman 2001; Pudlo et al. 2016). RF is a machine-learning algorithm that uses hundreds of bootstrapped decision trees to perform classification using a set of predictor variables, the summary statistics in this case. We grew a classification forest of 1000 trees based on all simulated datasets. After applying the random forest computation to the observed dataset, the scenario with the highest classification vote was selected as the most likely. Subsequently, we estimated its posterior probability using a second random forest procedure involving 1000 trees (Pudlo et al. 2016). To assess the overall performance of our ABC scenario selection, we (1) computed the prior error rate based on the available out-of-bag simulations, and (2) performed the scenario selection analysis using an alternative random selection of 10,000 synonymous SNPs, and a new reference table.

##### *2.3 Origin of the eastern North American population of* Harmonia axyridis*: results*

Our analysis suggested that scenario 1 was the most likely, indicating that the initial invasive population in North America originated from the eastern part of the Asian native area. This scenario received a total of 627 votes out of 1000, with a posterior probability of 0.764 (see table below). This conclusion was further supported by a low prior error rate (i.e. mean classification error) of 4.27%, strong robustness (Figure A2), and the analysis of the second set of SNPs which yielded 600 votes and a posterior probability of 0.720 for scenario 1 (see Table A1).

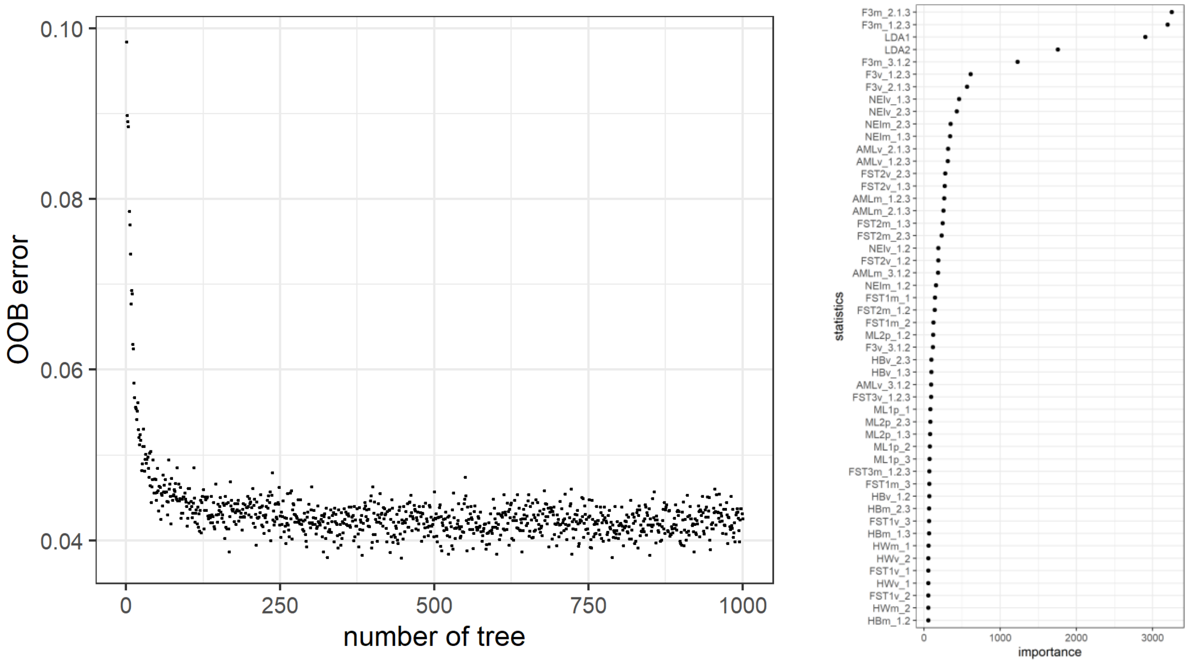


**Figure A2**: evaluation of the robustness of the ABC random forest analysis. Left panel: the evolution of the prior error rate (OOB error) relative to the number of trees in the forest shows that the choice of 1000 trees was sufficient. Right panel: the computation of variable importance using the Gini index showed that none of 5 noise variables added to the analysis were found among the 50 most informative statistics.

| Analysis | Votes scenario 1 | Votes scenario 2 | Votes scenario 3 | Selected scenario (post probability) |
|---|---|---|---|---|
| First dataset | 627 | 18 | 355 | 1 (0.764) |
| Second dataset | 600 | 41 | 359 | 1 (0.720) |

**Table A1**: ABC random forest votes of each scenario and posterior probability of the selected scenario regarding the origin of the eastern North American population of *Harmonia axyridis*.

To conclude, the use of a large set of genetic markers, comprising 10,000 biallelic SNPs, alongside an up-to-date machine learning algorithm, yielded a distinct result compared to the findings obtained with 18 microsatellites markers by Lombaert et al. (2011). We consider the result obtained in the current study as more robust, leading us to conclude that no admixture was responsible for the invasion of *Harmonia axyridis* in eastern North America.

#### 3. Bottleneck intensities for *Harmonia axyridis* and *Diabrotica virgifera virgifera*

##### *3.1 Background and motivation*

When a demographic bottleneck occurs in a population, it is likely to affect the genetic load. The nature of this impact theoretically depends on the intensity of the bottleneck (e.g., Glémin 2003), which results from a complex interaction between the number of founders and the duration of the bottleneck. Building on the above findings regarding the origin of the North-East American population of *Harmonia axyridis* and the abundant literature describing the invasion routes of *Harmonia axyridis* and *Diabrotica virgifera virgifera* (Miller et al. 2005; Ciosi et al. 2008; Lombaert et al. 2010, 2011, 2014, 2018), we used ABC analyses to simulate the final invasion scenarios of both species and to estimate the bottleneck intensities experienced by all invasive populations. While various parameter estimates are already available in the literature, this approach allowed us to estimate these parameters across both species within a unified framework.

##### *3.2 ABC analyses: simulating invasion scenario for* Diabrotica virgifera virgifera

For *Diabrotica virgifera virgifera*, the final invasion scenario was coded as follow (see Figure A3 for a graphical representation):

```
Nm Nc Np Ncse Ninw Ngh
0 sample 1
0 sample 2
0 sample 3
0 sample 4
0 sample 5
tinw-DBinw VarNe 5 NFinw
tinw merge 3 5
tcse-DBcse VarNe 4 NFcse
tcse merge 3 4
tp-DBp VarNe 3 NFp
tp merge 2 3
tc-DBc VarNe 2 NFc
tc VarNe 2 Nm
tgm merge 1 2
tm-DBn VarNe 1 NFm
tm merge 6 1
```

Population numbers are as follows: 1 for Mexico (effective population size $N_m$; sample D-N-MX1); 2 for Colorado (effective population size $N_c$; sample D-I-COL); 3 for Pennsylvania (effective population size $N_p$; sample D-I-PEN); 4 for Hungary (effective population size $N_{cse}$; sample D-I-HUN); 5 for North West Italy (effective population size $N_{nwi}$; sample D-I-ITA); 6 for an unsampled ancestral native population (effective population size $N_{gh}$). All *DB* and *NF* parameters represent, respectively, the bottleneck duration and the effective number of founders during this duration for the corresponding population. All t parameters represent the merging time of the corresponding population with its source population. In the case of Colorado, the source is an unsampled native population that merges into the native Mexican population at time $t_{gm}$. The native Mexican population, in turn, merges into the ancestral population at time $t_m$.

Historical and demographic parameter values for simulations were drawn from prior distributions defined as follows:

```
NFinw N LU[2,1000,0,0]
NFcse N LU[2,1000,0,0]
NFp N LU[2,1000,0,0]
NFc N LU[2,1000,0,0]
Nm N LU[100,100000,0,0]
NFm N LU[2,1000,0,0]
Nc N LU[100,100000,0,0]
Np N LU[100,100000,0,0]
Ncse N LU[100,100000,0,0]
Ninw N LU[100,100000,0,0]
Ngh N LU[100,100000,0,0]
tinw T UN[15,20,0,0]
DBinw T UN[1,10,0,0]
tcse T UN[23,28,0,0]
DBcse T UN[1,10,0,0]
tp T UN[30,35,0,0]
DBp T UN[1,10,0,0]
tc T LU[148,1500,0,0]
DBc T UN[1,10,0,0]
tgm T LU[148,1500,0,0]
tm T LU[148,1500,0,0]
DBn T UN[1,10,0,0]
```

With the following conditions:

```
tm>tgm
tgm>=tc
```

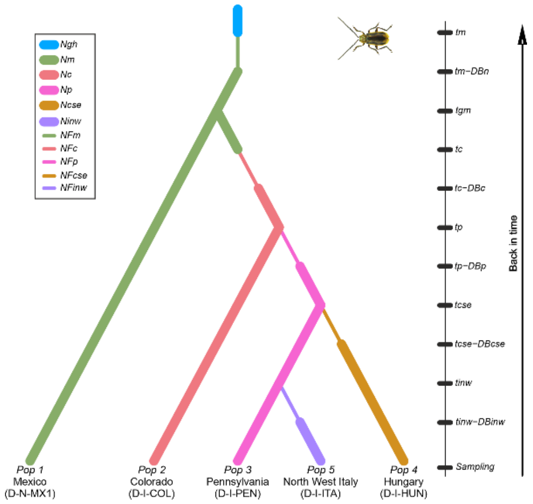


**Figure A3**: Schematic representation of the final invasion scenario for *Diabrotica virgifera virgifera* used in the ABC analyses to estimate the bottleneck intensity of each invasive population. Time is not to scale.

Since the dataset includes five populations, the total number of summary statistics was 292.

##### *3.3 ABC analyses: simulating invasion scenario for* Harmonia axyridis

For *Harmonia axyridis* (hereafter called HA), the final invasion scenario was coded as follow (see Figure A4 for a graphical representation):

```
Ne Nw Nlou Nwas Ngh
0 sample 1
0 sample 2
0 sample 3
0 sample 4
twas-DBwas VarNe 4 NFwas
twas VarNe 4 Ne
tlou-DBlou VarNe 3 NFlou
tlou VarNe 3 Ne
tgewas merge 1 4
tgelou merge 1 3
tne merge 5 1
tnw merge 5 2
```

Population numbers are as follows: 1 for East Asia (effective population size $N_e$; sample H-N-CHI); 2 for West Asia (effective population size $N_w$; sample H-N-RUS); 3 for eastern North America (effective population size $N_{lou}$; sample H-I-PEN); 4 for western North America (effective population size $N_{was}$; sample H-I-WAS); 5 for an unsampled ancestral native population (effective population size $N_{gh}$). All *DB* and *NF* parameters represent, respectively, the bottleneck duration and the effective number of founders during this duration for the corresponding population. All t parameters represent the merging time of the corresponding population with its source population. In the case of eastern and western North American populations, the sources are unsampled native populations that merge into the native east Asian population at the corresponding time $t_{ge}$. The native populations, in turn, merge into the ancestral population at the corresponding time $t_n$.

Historical and demographic parameter values for simulations were drawn from prior distributions defined as follows:

```
Ngh N LU[100,20000,0,0]
NFwas N LU[2,1000,0,0]
Ne N UN[100,100000,0,0]
NFlou N LU[2,1000,0,0]
Nw N UN[100,100000,0,0]
Nlou N UN[100,100000,0,0]
Nwas N UN[100,100000,0,0]
twas T UN[60,65,0,0]
DBwas T UN[1,10,0,0]
tlou T UN[68,73,0,0]
DBlou T UN[1,10,0,0]
tgewas T LU[60,3000,0,0]
tgelou T LU[68,3000,0,0]
tne T LU[100,3000,0,0]
tnw T LU[100,3000,0,0]
```

With the following conditions:

```
tgelou>=tlou
tne>=tgelou
tgewas>=twas
tne>=tgewas
```

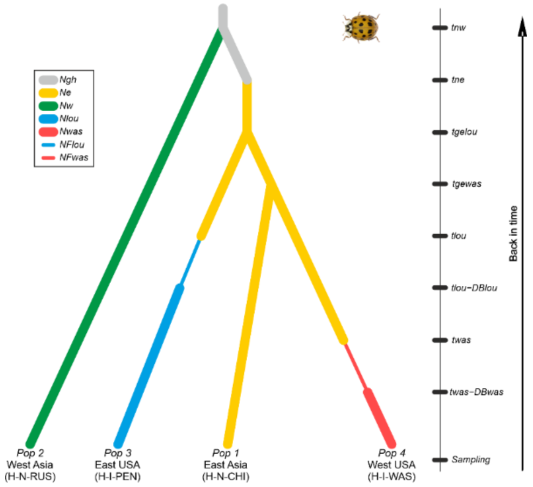


**Figure A4**: Schematic representation of the final invasion scenario for *Harmonia axyridis* used in the ABC analyses to estimate the bottleneck intensity of each invasive population. Thin lines indicate a bottleneck. Time is not to scale.

Since the dataset includes four populations, the total number of summary statistics was 130.

##### *3.4 ABC analyses: estimating bottleneck intensities*

For each scenario, we first assessed their goodness-of-fit by determining whether priors were off target, comparing the distributions of simulated summary statistics with the value of the observed dataset. Then, for each invasive population of both species, we estimated the composite parameter *DB/NF* using a regression by random forest methodology (Raynal et al. 2019), with classification forests consisting of 1,000 trees. This composite parameter allows us to measure the relative intensity of the bottleneck by incorporating both the effect of demography (*NF*, effective number of founder) and time (*DB*, bottleneck duration), which are intrinsically linked and whose effects on genetic diversity are difficult to disentangle.

##### 3.5 Estimation of bottleneck intensities: results

Comparisons of the distributions of simulated summary statistics with values from the observed dataset showed that the combination of scenarios and priors we chose was realistic. Indeed, only 17 out of 292 observed statistics (5.8%) for *Diabrotica virgifera virgifera* and 4 out of 130 observed statistics (3.1%) for *Harmonia axyridis* significantly (at a 5% threshold) lay in the tails of the probability distribution of statistics calculated from prior simulations.

Point estimates of composite parameter *DB/NF* (hereafter referred to as "bottleneck intensity") for all invasive populations of both species are presented in Figure A5. Overall, bottlenecks were found to be less severe in *Harmonia axyridis* (HA) than in *Diabrotica virgifera virgifera* (DVV). In DVV, only the Pennsylvania population (D-I-PEN) suffered a modest bottleneck, similar to those observed in the HA populations, which may be explained by its spatial connection to its source population in Colorado (Lombaert et al. 2018). The two European DVV populations show similar, relatively high bottleneck intensities. Finally, the oldest invasive DVV population in Colorado (D-I-COL) shows high estimates of bottleneck intensity, but with very large ranges, so these results should be interpreted with caution.

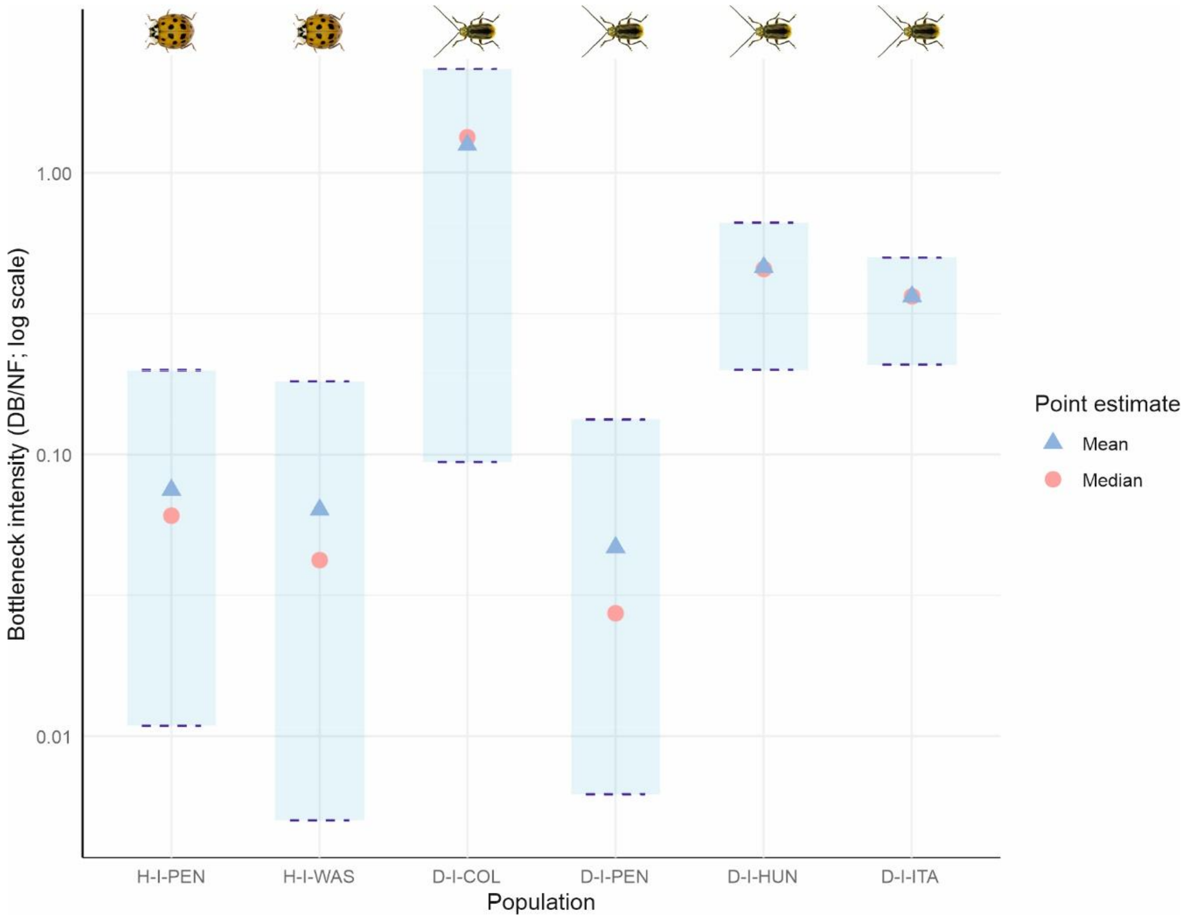


**Figure A5**: Estimation of the composite parameter *DB/NF* (bottleneck duration / effective number of founder) for each invasive population of *Harmonia axyridis* and *Diabrotica virgifera virgifera*. For each population, point estimates are the mean (blue triangle), the median (red circle), the 0.05 quantile (lower dash line) and the 0.95 quantile (upper dash line).

#### Appendix S2
#### Computation of the $R_{XY}$ statistic

The $R_{XY}$ statistic measures the relative excess or deficit of a specific category of variant (either missense or loss-of-function) in an invasive population (population *X*) compared to its source population (population *Y*). It is computed as described by Xue et al. (2015). For each category of non-synonymous mutation, we first calculate the following two ratios:

$$L_{X,Y}(C) = \frac{\sum_{i \in C} p_i^X q_i^Y}{\sum_{j \in S} p_j^X q_j^Y}$$

and:

$$L_{Y,X}(C) = \frac{\sum_{i \in C} p_i^Y q_i^X}{\sum_{j \in S} p_j^Y q_j^X}$$

Where *p* represents derived allele frequencies, *q* represents ancestral allele frequencies, *i* and *j* are nucleotide sites, *C* is the set of a specific category of non-synonymous protein-coding sites (either missense or loss-of-function), and *S* is the set of synonymous sites.

The ratio statistic is then defined as:

$$R_{X/Y}(C) = \frac{L_{X,Y}(C)}{L_{Y,X}(C)}$$

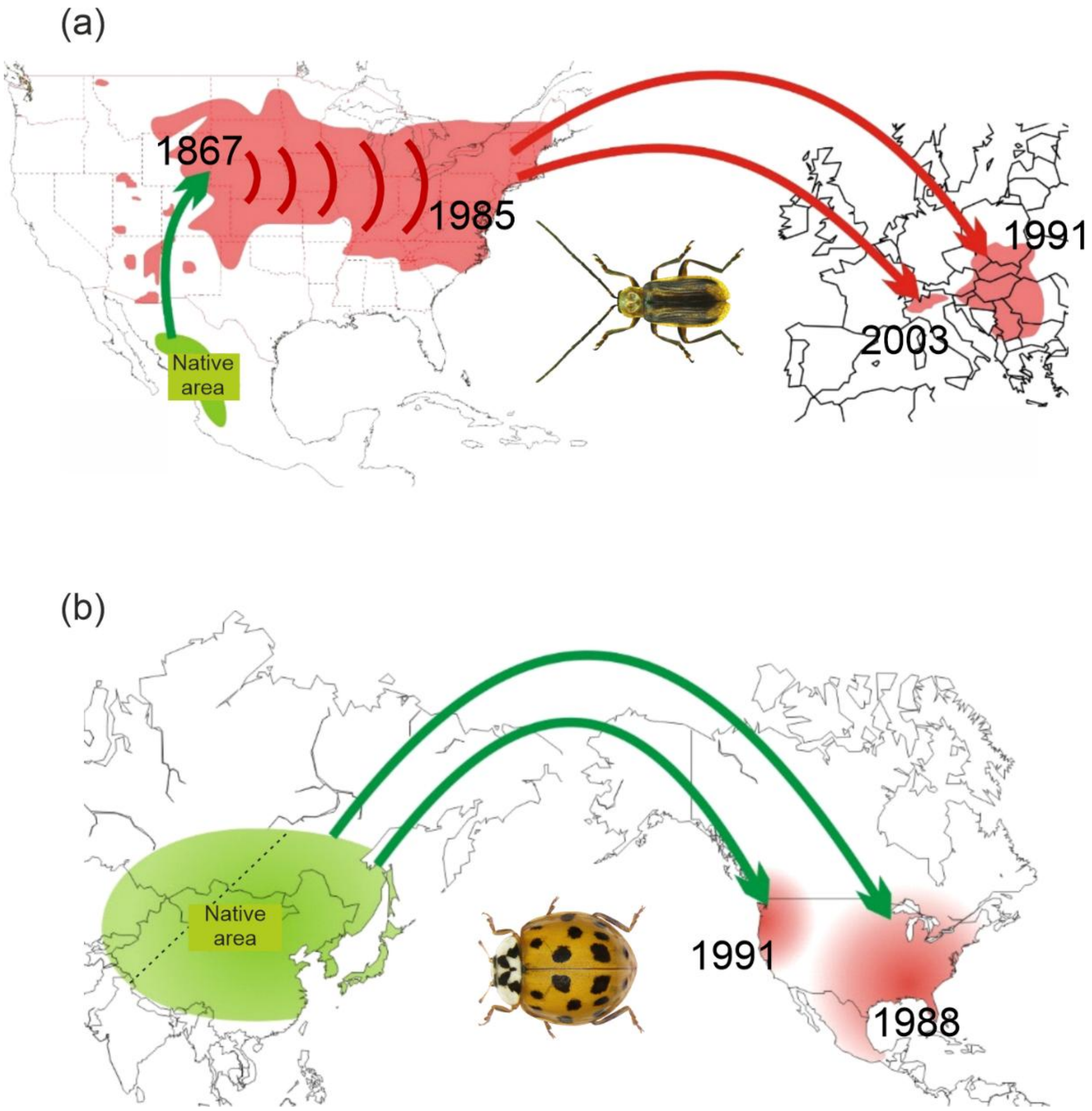


**Figure S1:** Known invasion routes of (a) *Diabrotica virgifera virgifera* and (b) *Harmonia axyridis*. Native areas are shown in green and invasive areas in red. The dashed line in the native range of *Harmonia axyridis* roughly delineates the western and eastern genetic clusters. Green arrows indicate introductions from a native population, while red arrows indicate introductions from an already invasive population. Years of first observation for invasive populations are indicated. Invasion routes were reconstructed from several previous studies (Miller et al. 2005; Ciosi et al. 2008; Lombaert et al. 2010, 2011, 2018), as well as the present study (see Appendix S1).

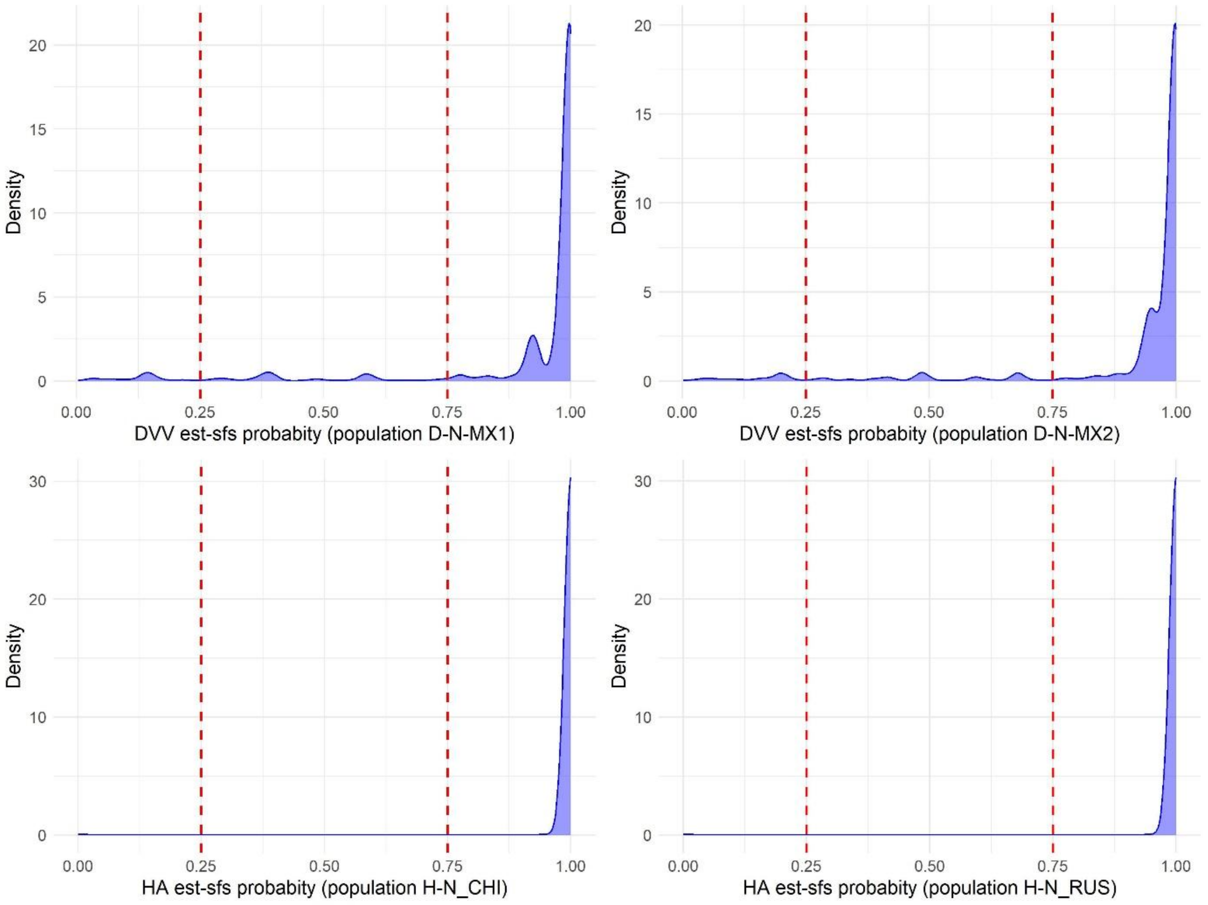


**Figure S2:** Distribution of est-sfs probabilities for each species and target population. The vertical dashed red lines illustrate the threshold applied in this study.

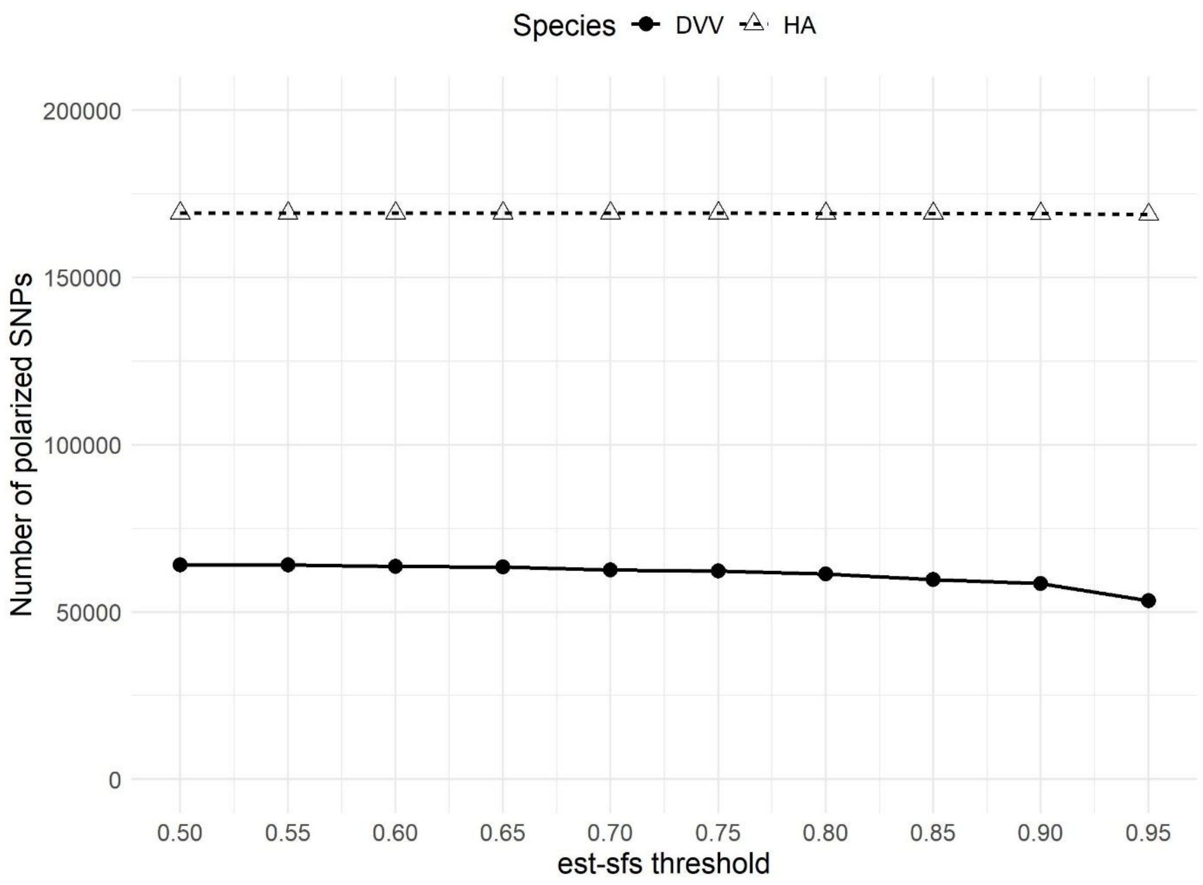


**Figure S3:** Number of polarized SNPs as a function of the est-sfs probability threshold.

(a)

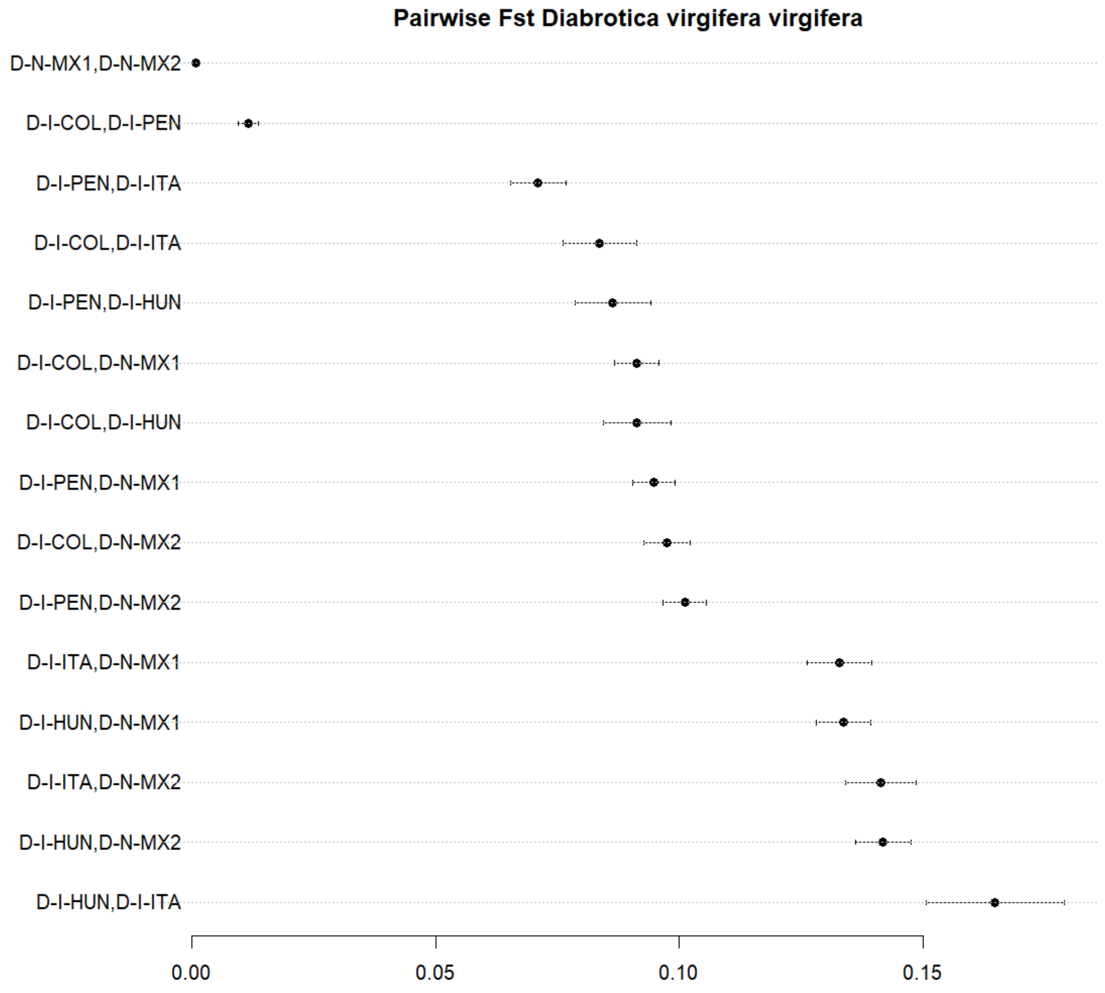


(b)

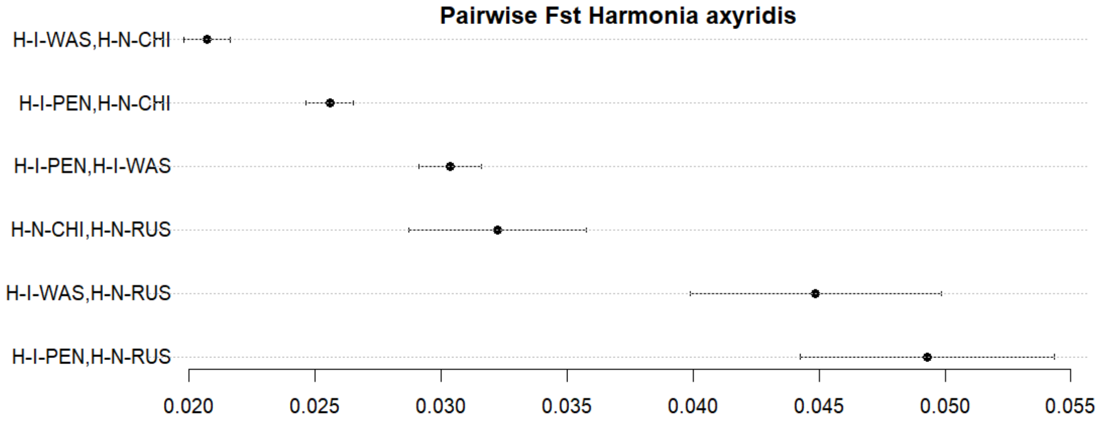


**Figure S4:** Pairwise $F_{ST}$ estimates between all sample pairs of (a) *Diabrotica virgifera virgifera* and (b) *Harmonia axyridis*. The error bars represent the 95% confidence interval computed from standard errors obtained through 100-block Jackknife resampling.

| Population code name | Species name | BioSample accession | SRA accession |
|---|---|---|---|
| D-N-MX1 | *Diabrotica virgifera virgifera* | SAMN40074446 | SRR28065669; SRR28065666; SRR28065667 |
| D-N-MX2 | *Diabrotica virgifera virgifera* | SAMN40074447 | SRR28065665; SRR28065663; SRR28065664 |
| D-I-COL | *Diabrotica virgifera virgifera* | SAMN40074444 | SRR28065674; SRR28065675; SRR28065673 |
| D-I-PEN | *Diabrotica virgifera virgifera* | SAMN40074445 | SRR28065671; SRR28065672; SRR28065670 |
| D-I-HUN | *Diabrotica virgifera virgifera* | SAMN40074443 | SRR28065677; SRR28065678; SRR28065676 |
| D-I-ITA | *Diabrotica virgifera virgifera* | SAMN40074442 | SRR28065652; SRR28065653; SRR28065654 |
| H-N-CHI | *Harmonia axyridis* | SAMN40074435 | SRR28065680 |
| H-N-RUS | *Harmonia axyridis* | SAMN40074436 | SRR28065679 |
| H-I-PEN | *Harmonia axyridis* | SAMN40074437 | SRR28065668 |
| H-I-WAS | *Harmonia axyridis* | SAMN40074438 | SRR28065658 |
| D-OG-ADE | *Diabrotica adelpha* | SAMN40074448 | SRR28065662 |
| D-OG-TRI | *Cerotoma trifurcata* | SAMN40074451 | SRR28065659 |
| H-OG-YED | *Harmonia yedoensis* | SAMN40074439 | SRR28065657 |
| H-OG-CON | *Harmonia conformis* | SAMN40074440 | SRR28065656 |
| H-OG-QUA | *Harmonia quadripunctata* | SAMN40074441 | SRR28065655 |

**Table S1:** Biosample and Sequence Read Archive (SRA) accessions for all samples used in this study (see Table 1 for additional information). All references can be found in public databases within BioProject PRJNA1079689: https://www.ncbi.nlm.nih.gov/bioproject/PRJNA1079689

| Sample | Capture 1 | Capture 2 | Capture 3 | Capture 4 | Capture 5 |
|---|---|---|---|---|---|
| D-N-MX1 | 250 | 0 | 250 | 250 | 0 |
| D-N-MX2 | 0 | 250 | 250 | 0 | 250 |
| D-I-COL | 250 | 0 | 250 | 250 | 0 |
| D-I-PEN | 0 | 250 | 250 | 0 | 250 |
| D-I-HUN | 250 | 250 | 0 | 0 | 250 |
| D-I-ITA | 250 | 250 | 0 | 250 | 0 |
| D-OG-ADE | 0 | 0 | 0 | 83.4 | 0 |
| D-OG-TRI | 0 | 0 | 0 | 83.4 | 0 |

**Table S2:** Quantity of amplified DNA (in nanograms) per population and outgroup samples within each *Diabrotica virgifera virgifera* exome capture reaction. See Table 1 for correspondence of sample names.

| Sample | Capture 1 | Capture 2 |
|---|---|---|
| H-N-CHI | 475 | 0 |
| H-N-RUS | 0 | 475 |
| H-I-PEN | 475 | 0 |
| H-I-WAS | 0 | 475 |
| H-OG-YED | 0 | 25 |
| H-OG-CON | 25 | 0 |
| H-OG-QUA | 25 | 0 |

**Table S3:** Quantity of amplified DNA (in nanograms) per population and outgroup samples within each *Harmonia axyridis* exome capture reaction. See Table 1 for correspondence of sample names.

| Species | Population code name | Total number of read pairs | Mbases | Number of cleaned read pairs | Number of cleaned reads (SE1 only) | Number of cleaned reads (SE2 only) | Total number of cleaned reads | Proportion of reads retained (%) |
|---|---|---|---|---|---|---|---|---|
| *Diabrotica virgifera virgifera* | D-N-MX1 | 44,817,432 | 13445.23 | 23,853,708 | 12,983,827 | 1,207,657 | 38,045,192 | 84.9% |
| *Diabrotica virgifera virgifera* | D-N-MX2 | 49,240,579 | 14772.17 | 27,135,813 | 13,481,989 | 1,498,006 | 42,115,808 | 85.5% |
| *Diabrotica virgifera virgifera* | D-I-COL | 46,195,866 | 13858.76 | 24,560,305 | 13,431,214 | 1,298,675 | 39,290,194 | 85.1% |
| *Diabrotica virgifera virgifera* | D-I-PEN | 44,388,533 | 13316.56 | 24,128,204 | 12,475,387 | 1,246,443 | 37,850,034 | 85.3% |
| *Diabrotica virgifera virgifera* | D-I-HUN | 60,055,627 | 18016.69 | 34,093,035 | 15,300,849 | 1,879,292 | 51,273,176 | 85.4% |
| *Diabrotica virgifera virgifera* | D-I-ITA | 56,482,967 | 16944.89 | 31,582,786 | 13,682,870 | 2,100,689 | 47,366,345 | 83.9% |
| *Harmonia axyridis* | H-N-CHI | 71,236,915 | 21371.07 | 48,980,633 | 16,721,947 | 1,528,100 | 67,230,680 | 94.4% |
| *Harmonia axyridis* | H-N-RUS | 62,668,899 | 18800.67 | 43,204,013 | 15,016,778 | 1,368,480 | 59,589,271 | 95.1% |
| *Harmonia axyridis* | H-I-PEN | 60,105,748 | 18031.72 | 39,582,696 | 8,821,330 | 6,341,663 | 54,745,689 | 91.1% |
| *Harmonia axyridis* | H-I-WAS | 59,431,143 | 17829.34 | 40,720,319 | 14,150,309 | 1,308,966 | 56,179,594 | 94.5% |
| *Diabrotica adelpha* | D-OG-ADE | 5,357,457 | 1607.24 | 3,921,123 | 1,254,138 | 33,513 | 5,208,774 | 97.2% |
| *Diabrotica balteata* | D-OG-BAL | 3,839,170 | 1151.75 | 2,887,561 | 818,348 | 27,020 | 3,732,929 | 97.2% |
| *Diabrotica undecimpunctata howardi* | D-OG-UND | 3,569,247 | 1070.77 | 2,857,981 | 597,593 | 43,982 | 3,499,556 | 98.0% |
| *Cerotoma trifurcata* | D-OG-TRI | 735,945 | 220.78 | 581,422 | 123,845 | 10,341 | 715,608 | 97.2% |
| *Harmonia yedoensis* | H-OG-YED | 2,008,136 | 602.44 | 1,576,084 | 392,346 | 14,198 | 1,982,628 | 98.7% |
| *Harmonia conformis* | H-OG-CON | 1,810,155 | 543.05 | 1,540,507 | 233,331 | 16,630 | 1,790,468 | 98.9% |
| *Harmonia quadripunctata* | H-OG-QUA | 1,102,318 | 330.69 | 961,755 | 121,459 | 9,229 | 1,092,443 | 99.1% |

**Table S4:** Summary statistics for the raw sequence data obtained for the 17 libraries (10 focal species and 7 outgroup species). SE1 and SE2 denote the first and second ends of paired-end reads, respectively. See Table S1 for BioSample and SRA accessions.

| Species | Raw vcf size after calling | Bi-allelic SNPs only | Filtered bi-allelic SNPs (polarized) | Coding regions only (polarized) | Synonymous (polarized) | Non synonymous (NS) | |
|---|---|---|---|---|---|---|---|
| | | | | | | NS missense (polarized) | NS LoF (polarized) |
| *Diabrotica v. virgifera* | 3,795,499 | 2,789,733 | 263,639 (254,309) | 66,274 (62,212) | 37,559 (34,164) | 27,808 (27,146) | 907 (902) |
| *Harmonia axyridis* | 4,044,301 | 2,647,316 | 411,679 (410,595) | 169,755 (169,223) | 116,190 (115,740) | 52,307 (52,226) | 1,258 (1,257) |

**Table S5:** Total number of SNPs called and retained at each stage of the pipeline for each species. Where applicable, the number of positions successfully polarized is shown in brackets. The last three columns correspond to SNPs from coding regions categorized by SNPEFF: synonymous, missense and loss-of-function (LoF) mutations.

#### Supporting information references


Beaumont MA, Zhang WY, Balding DJ (2002) Approximate Bayesian computation in population genetics. Genetics 162:2025–2035

Breiman L (2001) Random forests. Mach Learn 45:5–32. https://doi.org/10.1023/A:1010933404324

Chapin J, Brou V (1991) Harmonia axyridis (Pallas), the third species of the genus to be found in the United States (Coleoptera: Coccinellidae). Proc Entomol Soc Washingt 93:630–635

Choisy M, Franck P, Cornuet JM (2004) Estimating admixture proportions with microsatellites: comparison of methods based on simulated data. Mol Ecol 13:955–968. https://doi.org/10.1111/j.1365-294X.2004.02107.x

Ciosi M, Miller NJ, Kim KS, et al (2008) Invasion of Europe by the western corn rootworm, *Diabrotica virgifera virgifera*: Multiple transatlantic introductions with various reductions of genetic diversity. Mol Ecol 17:3614–3627. https://doi.org/10.1111/j.1365-294X.2008.03866.x

Collin F-D, Durif G, Raynal L, et al (2021) Extending Approximate Bayesian Computation with Supervised Machine Learning to infer demographic history from genetic polymorphisms using DIYABC Random Forest. Mol Ecol Resour 21:2598–2613

Glémin S (2003) How are deleterious mutations purged? Drift versus nonrandom mating. Evolution (N Y) 57:2678–2687. https://doi.org/10.1111/j.0014-3820.2003.tb01512.x

Hivert V, Leblois R, Petit EJ, et al (2018) Measuring genetic differentiation from Pool-seq data. Genetics 210:315–330. https://doi.org/10.1101/282400

Leblois R, Gautier M, Rohfritsch A, et al (2018) Deciphering the demographic history of allochronic differentiation in the pine processionary moth Thaumetopoea pityocampa. Mol Ecol 27:264–278. https://doi.org/10.1111/mec.14411

Lombaert E, Ciosi M, Miller NJ, et al (2018) Colonization history of the western corn rootworm (*Diabrotica virgifera virgifera*) in North America: insights from random forest ABC using microsatellite data. Biol Invasions 20:665–677. https://doi.org/https://doi.org/10.1007/s10530-017-1566-2

Lombaert E, Guillemaud T, Cornuet JM, et al (2010) Bridgehead effect in the worldwide invasion of the biocontrol harlequin ladybird. PLoS One 5:e9743. https://doi.org/e9743 10.1371/journal.pone.0009743

Lombaert E, Guillemaud T, Lundgren J, et al (2014) Complementarity of statistical treatments to reconstruct worldwide routes of invasion: the case of the Asian ladybird *Harmonia axyridis*. Mol Ecol 23:5979–5997. https://doi.org/10.1111/mec.12989

Lombaert E, Guillemaud T, Thomas CE, et al (2011) Inferring the origin of populations introduced from a genetically structured native range by approximate Bayesian computation: case study of the invasive ladybird *Harmonia axyridis*. Mol Ecol 20:4654–4670. https://doi.org/10.1111/j.1365-294X.2011.05322.x

Miller N, Estoup A, Toepfer S, et al (2005) Multiple transatlantic introductions of the western corn rootworm. Science (80- ) 310:992. https://doi.org/10.1126/science.1115871

Nei M (1972) Genetic Distance between Populations. Am Nat 106:

Patterson N, Moorjani P, Luo Y, et al (2012) Ancient admixture in human history. Genetics 192:1065–1093. https://doi.org/10.1534/genetics.112.145037

Pudlo P, Marin JM, Estoup A, et al (2016) Reliable ABC model choice via random forests. Bioinformatics 32:859–866. https://doi.org/10.1093/bioinformatics/btv684

Raynal L, Marin J-M, Pudlo P, et al (2019) ABC random forests for Bayesian parameter inference. Bioinformatics 35:1720–1728. https://doi.org/10.24072/pci.evolbiol.100036

Weir BS, Goudet J (2017) A Unified Characterization of Population Structure and Relatedness. Genetics 206:2085–2103

Xue Y, Prado-martinez J, Sudmant PH, et al (2015) Mountain gorilla genomes reveal the impact of long-term population decline and inbreeding. Science (80- ) 348:242–245. https://doi.org/10.1126/science.aaa4484
